# iSTTC: a robust method for accurate estimation of intrinsic neural timescales from single-unit recordings

**DOI:** 10.1101/2025.08.01.668071

**Authors:** Irina Pochinok, Ileana L. Hanganu-Opatz, Mattia Chini

## Abstract

Intrinsic neural timescales (ITs) are an emerging measure of how neural circuits integrate information over time. ITs are dynamically regulated by behavioral context and cognitive demands, making them suitable for mapping high-level cognitive phenomena onto the underlying neural computations. In particular, IT measurements derived from single-unit activity (SUA) offer fine-grained resolution, critical for mechanistically linking individual neuron dynamics to cognition. However, current methods for estimating ITs from SUA suffer significant biases and instabilities, particularly when applied to sparse, noisy, or epoched neural spike data. Here, we introduce the intrinsic Spike Time Tiling Coefficient (iSTTC), a novel metric specifically developed to address these limitations. Leveraging synthetic and experimental single-unit recordings, we systematically assessed the performance of iSTTC relative to traditional approaches. Our findings demonstrate that iSTTC provides more accurate estimates of neural timescales across a wide range of conditions, reducing estimation error especially under challenging yet biologically relevant conditions. Crucially, iSTTC can be applied to both continuous and epoched data, overcoming a critical limitation of existing methods. Furthermore, iSTTC substantially relaxes inclusion criteria, increasing the fraction of neurons suitable for analysis and thereby improving the representativeness and robustness of IT measurements. The methodological advances introduced by iSTTC represent a substantial step forward in accurately capturing neural circuit dynamics, ultimately enhancing our ability to link neural mechanisms to cognitive phenomena.

## 1 Introduction

A remarkable ability of our brains is that of being capable of choosing an appropriate course of action over extremely different timescales: from a rapid millisecond-range reaction to avoid an unforeseen obstacle while driving a car, to choosing the next move in a complex chess match, which might unfold over several minutes. Intrinsic neural timescales (ITs) are an emerging fundamental measure quantifying how different systems integrate information over time [1, 2]. ITs are defined as the decay rate of the auto-correlation of neural signals, and vary significantly across the brain [1]. This is thought of reflecting the computational properties of different brain regions. Within the neocortex, regions that sit at the bottom of the cortical hierarchy, such as primary sensory cortices, typically exhibit short timescales, conducive to rapid sensory processing. Higher-order brain areas, such as the prefrontal cortex, display substantially longer timescales, better suited to supporting more complex cognitive functions [1, 3]. Remarkably, these findings have been robustly replicated across phylogenetically distant species and disparate recording techniques, including single-unit activity (SUA), 2-photon and widefield calcium imaging, local field potentials (LFP), electrocorticography (ECoG), electro- and magneto-encephalography (EEG and MEG), and functional magnetic resonance imaging (fMRI) [2, 4, 5, 3, 6, 7, 8, 9, 10, 11, 12, 13, 14].

While few experimental studies have mechanistically investigated the factors that underlie this IT heterogeneity, early indications point to several single-neuron and network-level factors as being potentially implicated [3, 15, 16, 17, 18]. For instance, higher proportions of somatostatin-expressing interneurons and prolonged NMDA currents positively correlate with longer ITs, whereas parvalbumin-expressing interneurons generally contribute to shorter ITs [3, 16]. Importantly, ITs are not fixed but dynamically modulated by behavioral context, cognitive demands, and task engagement [10, 3, 6, 9, 19]. Another important property of ITs lies in their predictive power for behavioral and cognitive performance. For example, shorter ITs correlate with faster reaction times [20]. This suggests that modulation of ITs could be a mechanism by which neural circuits dynamically adjust computational resources to behavioral needs. Thus, a detailed understanding of the mechanisms regulating ITs might represent a promising approach to bridge neural dynamics with higher-order cognitive functions, and could even serve as mechanistic biomarkers in clinical contexts [21, 22, 23]. To this aim, recording modalities that bridge single neuron and network levels of investigations, such as SUA and 2-photon calcium imaging, are particularly promising for advancing our mechanistic understanding of ITs.

Despite their potential importance, current methods for estimating ITs from SUA exhibit substantial limitations [24]. Typically, IT estimation involves a two-step process: first calculating an autocorrelation (or autocorrelation-like) function (ACF, but see below), and subsequently fitting an exponential decay function [24]. Of note, in this paper we reserve the use of “ACF” for the classically defined autocorrelation function. Any alternative preprocessing, smoothing, or surrogate-based approximations are referred to as “ACF-like” methods. Recent methodological improvements have primarily focused on optimizing the exponential fitting step, with approaches such as adaptive Approximate Bayesian Computation (aABC) substantially reducing bias and improving robustness [25]. However, significant uncertainty remains in the initial process of estimating the ACF itself. This is a critical step whose results are influenced by multiple factors, including sensitivity to data length [26], the SUA firing rate, the eventual presence of trials, the representation of the spiking data as binary or continuous, the choice of bin width or smoothing kernel, and other preprocessing choices. Furthermore, most existing trial-based methods impose strict inclusion criteria, thus restricting analysis to a minority of recorded neurons [1, 27, 7, 28, 29, 30]. This runs counterintuitive to the notion that ITs are based on the decay rate of all neurons in a certain brain area. Additionally, given that the ACF cannot be readily estimated on epoched, non-continuous data, the most commonly used approach for trial-based IT estimation (referred to as PearsonR in this manuscript, see Materials and Methods for details) does not calculate the genuine ACF, but instead measures pseudo-autocorrelations on trial-averaged data [1].

To address these limitations, we introduce a novel metric: the intrinsic Spike Time Tiling Coefficient (iSTTC). iSTTC, an extension of STTC [31], robustly estimates ITs and offers significant improvements over traditional ACF-based approaches. It provides more accurate IT estimates on long, uninterrupted spiking data, particularly under biologically realistic cortical conditions, characterized by low firing rates and high recurrent connectivity. Unlike traditional ACF methods, iSTTC is unaffected by zero-padding, and thus also seamlessly generalizes to epoched spiking data after 0-padding and concatenation. In simulated trial-based data, iSTTC consistently outperforms the PearsonR method, producing more accurate estimates of the true underlying timescales, particularly if the number of trials is low. Lastly, iSTTC relaxes the stringent inclusion criteria required by other commonly used trial-based methods, thus enabling IT estimation in a substantially larger fraction of neurons. This broader applicability enhances both the statistical power and representativeness of the estimated ITs.

In summary, iSTTC is a robust, accurate, and broadly applicable tool for IT estimation from SUA signals. By improving the reliability of IT measurements, iSTTC advances our ability to link intrinsic neural dynamics to functional and behavioral outcomes, strengthening ITs’ potential role as mechanistic biomarkers for cognitive processing.

## 2 Results

### 2.1 iSTTC calculates an autocorrelation-like function on non-binned spike trains

ITs are defined as the decay time constant of the signal’s autocorrelation function (Figure 1A). Estimation of ITs is a two-step process (Figure 1B): first, computing the ACF (or ACF-like), and second, estimating its decay time constant. Methods for estimating decay time constant based on the ACF/ACF-like can be broadly categorized into model-free and model-based approaches [24]. In this paper we focus on optimizing the first step (the calculation of the ACF/ACF-like) and use a model-based approach that assumes that the ACF/ACF-like follows an exponential decay (Figure 1C).

**Figure 1:**
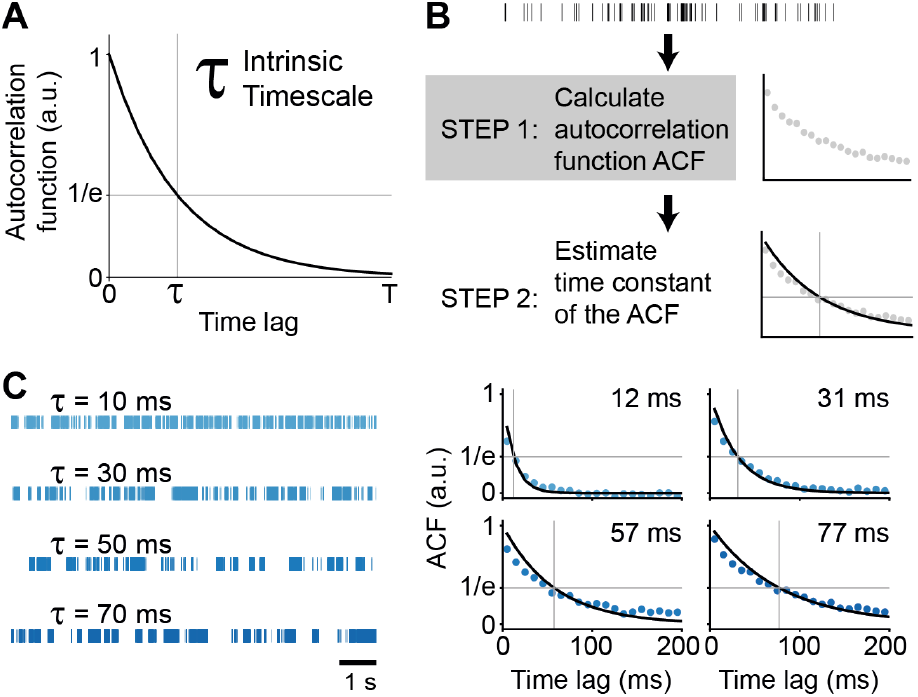
Definition and schematic representation of intrinsic timescale estimation. **(A)** Schematic illustration of an intrinsic timescale, defined as the decay time constant of the spike train ACF. **(B)** Schematic of the intrinsic timescale estimation pipeline for single-unit activity. **(C)** Representative examples of spike trains with known and increasing from top to bottom intrinsic timescales (left) and their corresponding estimated intrinsic timescales (right). In **(C)**, blue dots represent the computed ACF, and the black line indicates the fitted exponential decay function.

The calculation of the ACF/ACF-like, although seemingly straightforward, can be influenced by several arbitrary choices. First, the time series used to estimate the ITs can differ in structure: they may consist of long continuous spike trains [32] or a set of shorter trials, typically taken from baseline or fixation periods within behavioral tasks when the subject is not actively engaged [1, 27]. Second, spiking data can be represented either as binary spike times or as continuous signals obtained by binning or smoothing the spike train, which influences the method used to calculate the ACF/ACF-like. Both the structure of the data and the signal representation can introduce variability in the resulting ACF/ACF-like and, by extension, in the IT estimates.

To investigate the effects of these factors, we estimated ITs on a large synthetic dataset and an experimental one (Table 1).

**Table 1:** Summary of used datasets.

|  | **Synthetic SUA** | **Mouse single unit activity** |
| --- | --- | --- |
| **Reference** | Datasets were generated in this study | [2] |
| **Dataset source** | gin.g-node.org/iinnpp/isttc | Openly available for download in Neurodata Without Borders format via the AllenSDK |
| **Dataset info** | 10<sup>5</sup> single units with varying spike train features (firing rate, intrinsic timescale and burstiness) | 5775 single units, 26 mice, 8 brain areas, 30 min long spike trains and trials |
| **Related figures** | Figure 2, Figure 3, Figure 4, Figure 5 | Figure 6 |
| **Investigated biases** | Trial number, signal length, continuous vs epoched data, firing rate, IT magnitude, excitation strength | Trial number, signal length, continuous vs epoched data |

To calculate the ACF/ACF-like, we used three different methods (Table 2). Two of these are commonly used in the literature: i) the classic ACF for continuous signals; ii) the Pearson’s correlation coefficient on averaged signal (PearsonR) for epoched data. We compared these methods to iSTTC, the measure that we introduce in this manuscript (Supp. Figure 1A-B). iSTTC can be used as an ACF-like measure for both continuous and epoched data.

**Table 2:** Summary of used methods.

|  | **Binary signal**<br>**(spike times)** | **Binned spike**<br>**train** |
| --- | --- | --- |
| **Continuous signal** | iSTTC | ACF |
| **Trials** | iSTTC (trials) | PearsonR |

To compute an ACF-like of continuous signals using iSTTC, we generate lagged versions of the spike train by shifting it in time. For each lag *t*, we truncate the shifted train A by removing spikes occurring before *t*, and truncate the unshifted train B by removing spikes occurring at or beyond the signal length minus *t*. This alignment ensures both spike trains remain of equal length. We then compute the classic STTC between these two aligned trains, producing an ACF-like across lags (e.g., 50 ms, 100 ms, etc.) (Supp. Figure 1B) (see Materials and Methods for details).

For epoched data, before applying iSTTC, we linearize the signal by concatenating the epoched data, with zero-padded intervals inserted between trials. The ACF-like is then computed in the same way as for continuous spike data. By design, zero padding does not affect the iSTTC value (see Materials and Methods for details).

### 2.2 iSTTC outperforms ACF in IT estimation accuracy on continuous spiking data

To compare the performance of iSTTC and ACF in estimating ITs on long, continuous spiking data, we used a self-exciting Hawkes point process to simulate 10^5^ spike trains of 10 minutes each (Figure 2A, left). The spike trains varied systematically in time constant of the Hawkes point process (i.e. the IT to estimate; from 50 to 300 ms), firing rate (from 0.01 to 10 Hz), and excitation strength (from 0.1 to 0.9; see Materials and Methods for details). Each parameter was uniformly sampled within the specified range (Figure 2A, right). To compare the IT estimation accuracy of the two methods, for each iteration we computed the Relative Estimation Error (REE), defined as 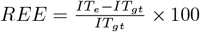, where *IT*_*e*_ is the estimated IT and *IT*_*gt*_ is the ground truth IT (Figure 2B).

**Figure 2:**
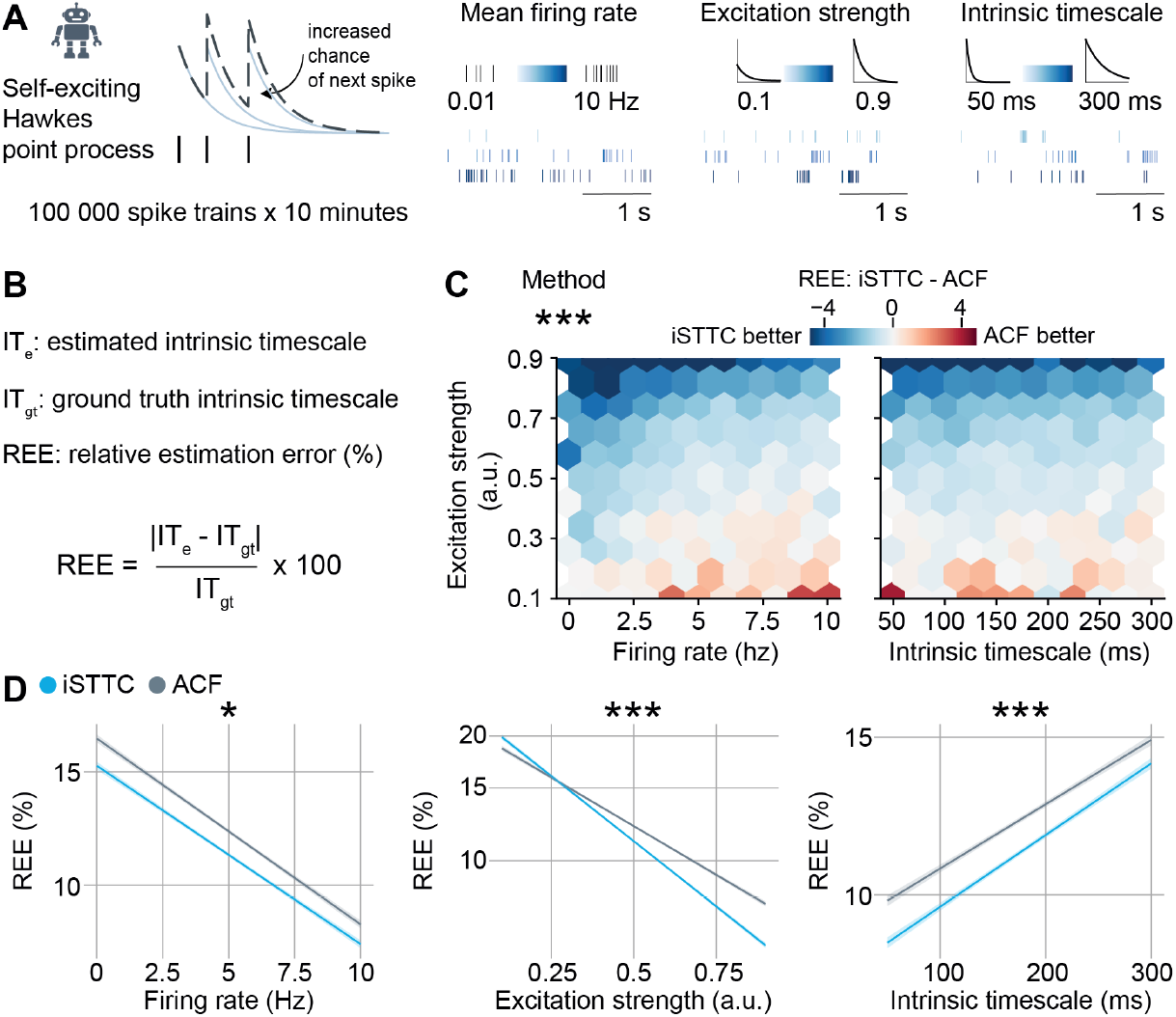
iSTTC is a better IT estimator than ACF on continuous data, particularly for low-firing rate and bursty units. **(A)** Schematic illustration of the synthetic dataset generation (left), and the underlying parameters with corresponding representative spike train examples (right). **(B)** Definition of the relative estimation error (REE) metric. **(C)** Hexbin plot displaying the difference in REE between iSTTC and ACF method as a function of firing rate and excitation strength (left), and IT and excitation strength (right) (n = 10^5^ single units). Color codes for the median REE difference in each bin, with blue indicating better IT estimation for iSTTC. **(D)** Line plot displaying predicted REE values for iSTTC and ACF as a function of firing rate (left), excitation strength (middle), and IT (right) (n = 10^5^ single units). Shaded areas represent 95% confidence intervals. Y-axes are plotted on a *log*_10_ scale. In **(C)**, asterisks indicate a significant effect of the IT estimation method. In **(D)**, asterisks indicate a significant effect of an interaction between method and firing rate (left), method and excitation strength (middle), and method and IT (right). * *p <* 0.05., *** *p <* 0.001. Generalized linear model with interactions **(C), (D)**.

Across the overwhelming majority of the explored parameter space, iSTTC yielded lower REEs than ACF (Figure 2C). To rigorously quantify this difference, we modeled how the IT estimation method impacted REEs with a multivariate generalized linear model. Across the entirety of the synthetic dataset, we found a robust effect of IT estimation method (coefficient estimate = –0.037, 95% CI [−0.039, −0.034], p *<* 10^*−*16^; Supp. Figure 2A), indicating that the REE of iSTTC was, on average, ~8% lower than that of ACF. This difference was not uniform throughout the parameter space (Figure 2C). Accordingly, statistical modeling revealed that the effect of IT estimation method significantly interacted with firing rate (coefficient estimate = 0.003, 95% CI [0.0007, 0.0061], p = 0.0124; Figure 2D, left; Supp. Figure 2A), excitation strength (coefficient estimate = −0.037, 95% CI [−0.039, −0.034], p *<* 10^*−*16^; Figure 2D, middle; Supp. Figure 2A) and the value of the IT (coefficient estimate = 0.005, 95% CI [0.003, 0.009], p *<* 10^*−*6^; Figure 2D, right; Supp. Figure 2A). Thus, estimating ITs with iSTTC resulted in lower REEs than ACF, an advantage that was particularly evident in low-firing rate, high excitation strength and low IT values regimes (Figure 2C-D).

While firing rate and IT are parameters that can be evaluated also in experimental datasets, excitation strength is not. Thus, we set out to investigate whether we could identify an experimentally-observable proxy for excitation strength, and reasoned that it might be reflected in the burstiness of the spike trains. To this aim, we quantified the local variation (Lv) [33], a measure of the regularity of the temporal spiking dynamics (Supp. Figure 2B). This coefficient takes values of 1 if the unit has random spiking dynamics; below 1 if the unit has regular spiking (i.e. oscillatory); and above 1 if the unit is bursty (Supp. Figure 2B). In line with our intuition, the Lv very tightly correlated with excitation strength (coefficient estimate = 0.436, 95% CI [0.434, 0.439], p *<* 10^*−*16^; Supp. Figure 2C, 2D, left) and thus also significantly interacted with the IT estimation method effect on REE (coefficient estimate = −0.042, 95% CI [−0.046, −0.039], p *<* 10^*−*16^; Supp. Figure 2D, right). Perhaps more importantly, this analysis revealed that the regimes in which ACF is a better IT estimator than iSTTC are regimes in which Lv *<* 1 (crossover point = 1.099), the units are regular-spiking (Supp. Figure 2D, right), and thus do not display the exponentially decaying autocorrelation that is a prerequisite to compute an IT.

Taken together, this data demonstrates that, on continuous spiking data, iSTTC is a better IT estimator than ACF, particularly in regimes that resemble neocortical brain activity, and are characterized by low and bursty firing.

### 2.3 iSTTC outperforms PearsonR in IT estimation accuracy on epoched spiking data

A key advantage of iSTTC over ACF is that the former is insensitive to 0-padding. Consequently, if the data on which IT estimation is based is epoched, as commonly observed in much of the current IT literature [1, 34, 7, 5, 27], iSTTC can still be readily employed after 0-padding and concatenating the epoched spiking data. Given that ACF cannot be used on epoched data, we compared iSTTC to the PearsonR approach that is commonly used in the literature [1, 34, 27]. To this aim, we used the same dataset of 10^5^ spike trains that we previously described. We extracted pseudo trials by randomly sampling 40 1s long chunks of spiking data (Figure 3A). The choice of trial number and length was based on values commonly used in the literature [1, 27, 34, 7]. Similarly to our approach in the previous section, we compared the IT estimation accuracy of the two methods by computing the REE for each iteration, and modeled the results with a generalized linear model.

**Figure 3:**
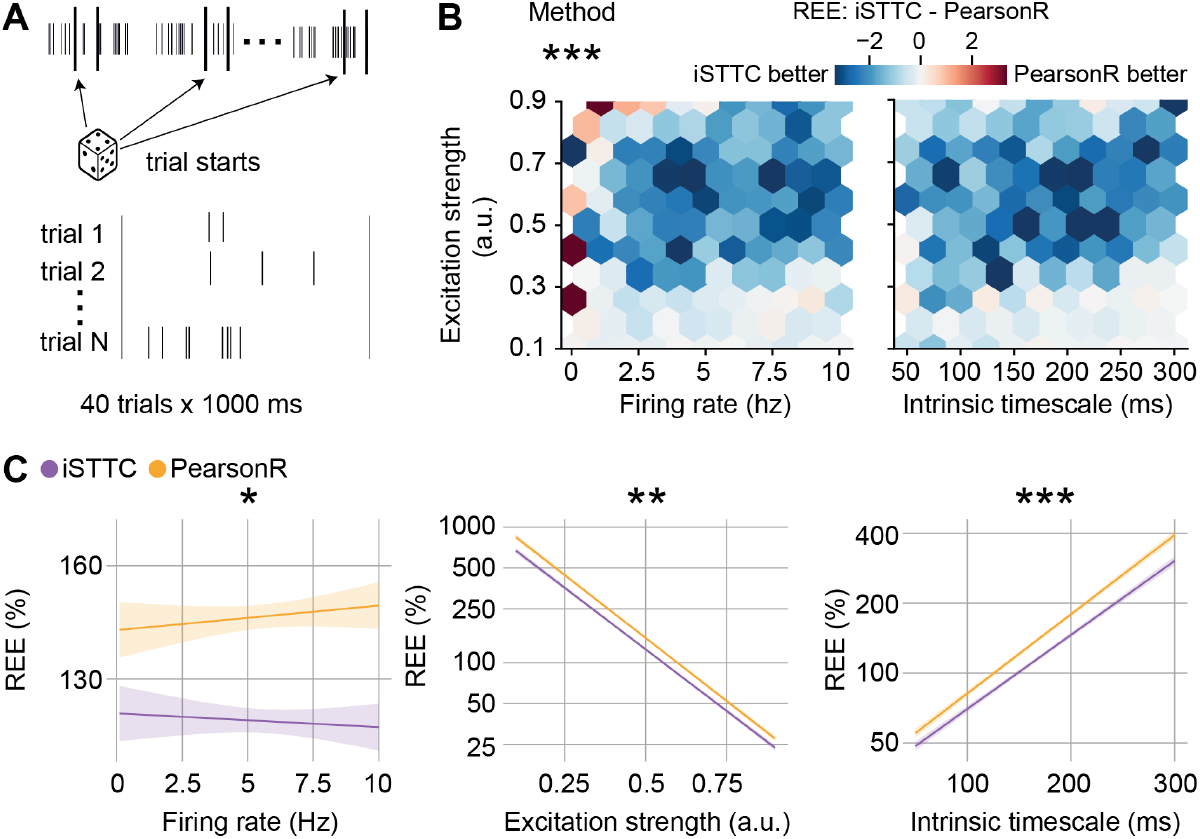
iSTTC provides better IT estimates from epoched spiking activity. **(A)** Schematic illustration of the generation of epoched data based on randomly sampled continuous spike trains. **(B)** Hexbin plot displaying the difference in REE between iSTTC and PearsonR as a function of firing rate and excitation strength (left), and IT and excitation strength (right) (n = 10^5^ single units, 40 trials x 1000 ms each). Color codes for the median REE difference in each bin, with blue indicating lower error for iSTTC. **(C)** Line plot displaying predicted REE values for iSTTC and PearsonR as a function of firing rate (left), excitation strength (middle), and IT (right) (n = 10^5^ single units, 40 trials x 1000 ms each). Shaded areas represent 95% confidence intervals. Y-axes are plotted on a *log*_10_ scale. In **(B)**, asterisks indicate a significant effect of the method. In **(C)**, asterisks indicate a significant effect of an interaction between method and firing rate (left), method and excitation strength (middle), and method and IT (right). * *p <* 0.05., ** *p <* 0.01., *** *p <* 0.001. Generalized linear model with interactions **(B), (C)**.

Across the overwhelming majority of the explored parameter space, iSTTC yielded lower REEs than ACF (coefficient estimate = −0.083, 95% CI [−0.090, −0.077], p *<* 10^*−*16^; Figure 3B, Supp. Figure 3A). On average, iSTTC estimates were 17.5% better than PearsonR (Supp. Figure 3A). This difference was not uniform throughout the parameter space (Figure 3B). Accordingly, the effect of IT estimation method significantly interacted with firing rate (coefficient estimate = −0.008, 95% CI [−0.014, −0.001], p = 0.02124; Figure 3C, left; Supp. Figure 3A), excitation strength (coefficient estimate = 0.009, 95% CI [0.002, 0.016], p = 0.00985; Figure 3C, middle; Supp. Figure 3A) and the value of the IT (coefficient estimate = −0.015, 95% CI [−0.023, −0.009], p *<* 10^*−*6^; Figure 3C, right; Supp. Figure 3A). Thus, estimating ITs with iSTTC results in lower REEs than PearsonR, an advantage that is particularly evident in high firing rate, low excitation strength and high IT values regimes (Figure 3B-C; Supp. Figure 3A). Notably, the magnitude of these interaction effects was much more modest than in the continuous spiking dataset (compare Supp. Figure 2A and Supp. Figure 3A). Perhaps most importantly, it is noteworthy that, regardless of the IT estimator that was employed, the REEs obtained from epoched data are roughly one order of magnitude larger than those obtained from continuous spiking data.

Taken together, this data indicates that, after 0-padding and trial concatenation, iSTTC outperforms PearsonR in estimating IT on epoched data. Moreover, the large REEs obtained from epoched data warrant some precaution in interpreting IT estimates obtained through this approach.

### 2.4 Epoched spiking data leads to vastly larger REEs than continuous spiking data

Inspired by the large difference on REEs between continuous and epoched data, we decided to systematically investigate how the accuracy of IT estimations depends on the amount and type of data that it is based upon. We first compared the REE obtained from the same spiking units in the continuous and epoched condition. To create the epoched dataset, we generated pseudo-trials by randomly selecting 40 1s segments of spiking activity for each unit. We confirmed our prior results about epoched data resulting in vastly larger REEs (Figure 4A, left). Irrespective of the IT estimation method, the epoched spiking data yielded a ~20% decrease of IT estimates with an REE below 100%, and a ~60% decrease of estimates with an REEs below 50% (Figure 4A, middle). In the aggregate, REEs from epoched data were an order of magnitude larger than continuous data (Figure 4A, right). Additionally, this analysis confirmed that iSTTC outperforms ACF and PearsonR on continuous and epoched data, respectively (Figure 4A). However, the advantage conferred by iSTTC over PearsonR is of limited magnitude when compared to the increase of REE due to epoched spiking data.

**Figure 4:**
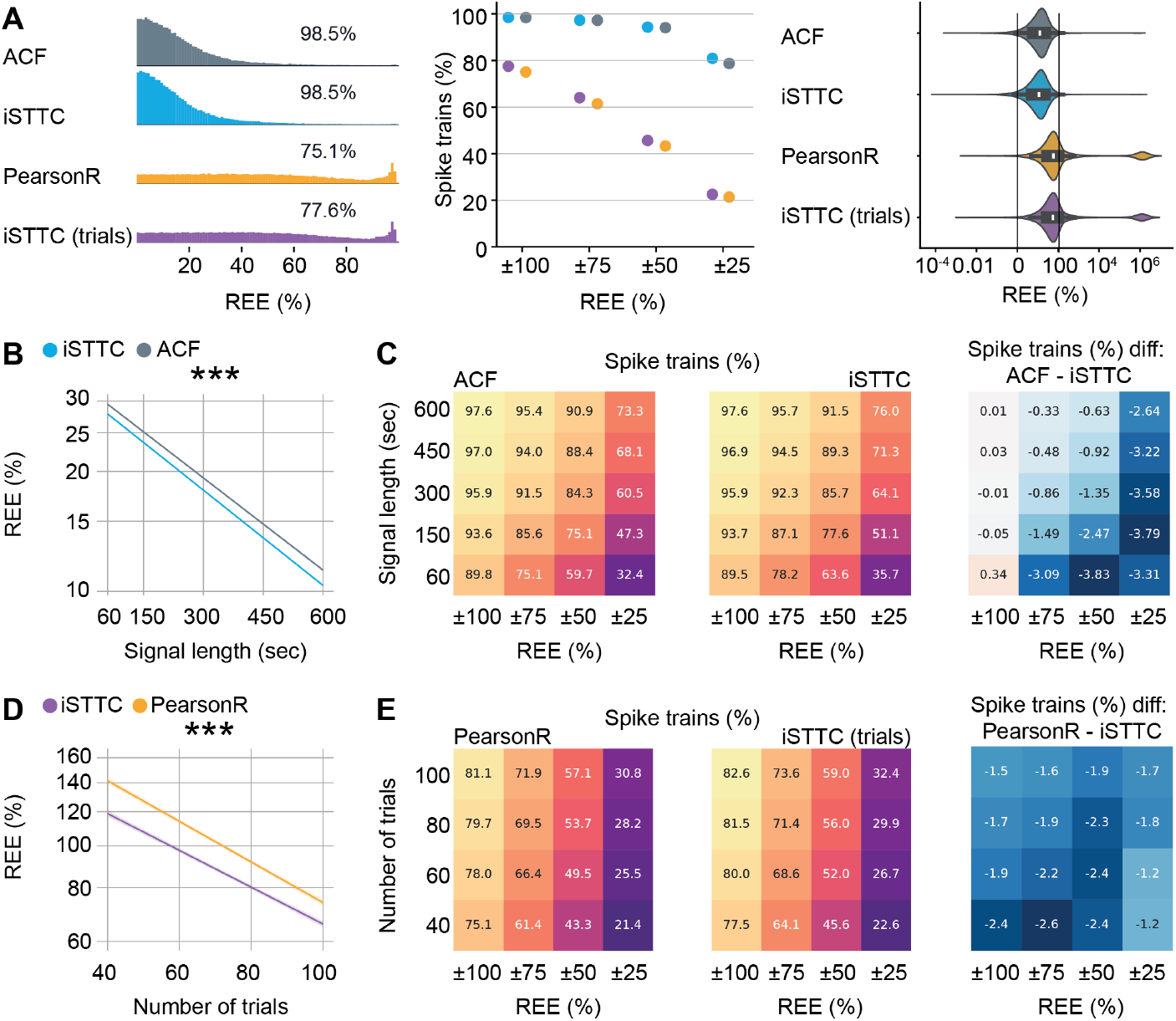
Relative estimation errors are higher for epoched than continuous spiking data. **(A)** Comparison of IT estimation accuracy across four methods. Ridgeline plot displaying the distribution of REE. Only spike trains with REE between 0% and 100% are shown. Percentages indicate the proportion of spike trains within this interval for each method (left). Scatter plot displaying the percentage of spike trains with REE falling within progressively narrower intervals (middle). Violin plot displaying the full distribution of REE values for each method. (right). **(B)** Line plot displaying predicted REE values for ACF and iSTTC as a function of signal length (n = 10^5^ single units per signal length). Shaded areas represent 95% confidence intervals. Y-axes are plotted on a *log*_10_ scale. **(C)** Heatmap displaying the percentage of spike trains with REE within specific intervals for ACF (left) and iSTTC (middle) across varying signal lengths. Color codes for the proportion of spike trains, with warmer colors indicating higher percentages of spike trains. (Right) Heatmap displaying the difference in performance between methods, computed as the difference between ACF and iSTTC. Negative values indicate better performance (lower REE) for iSTTC. Color codes for the magnitude of the difference in percentages. **(D)** Same as **(B)** for PearsonR and iSTTC. **(E)** Same as **(C)** for PearsonR and iSTTC. In (A) right, data is presented as median, 25th, 75th percentile, and interquartile range, with the shaded area representing the probability density distribution of the variable. In **(B)**, asterisks indicate a significant effect of interaction between method and signal length. In **(D)**, asterisks indicate a significant effect of an interaction between method and the number of trials. *** *p <* 0.001. Generalized linear model with interactions **(B), (D)**.

To further explore how the amount of data affects the REE of different methods, we systematically explored the effect of varying the amount of continuous (from 60s to 10 minutes) data and the number of trials (from 40 to 100 trials) on which we estimated ITs.

Increasing the amount of continuous spiking data led to a log-linear reduction in the REE of iSTTC and ACF alike (Figure 4B). Across the entirety of the parameter space, iSTTC outperformed ACF by an average of ~7% (coefficient estimate = −0.031, 95% CI [−0.033, −0.029], p *<* 10^*−*16^; Supp. Figure 4A) and consistently displayed a higher proportion of IT estimates within various ranges of the actual IT value (Figure 4C). The effect of IT estimation method interacted with the spike train length, firing rate, excitation strength and the IT value (signal length: coefficient estimate = −0.005, 95% CI [−0.152, −0.150], p *<* 10^*−*11^; firing rate: coefficient estimate = 0.002, 95% CI [0.0003, 0.0034], p = 0.017; excitation strength: coefficient estimate = −0.028, 95% CI [−0.030, −0.027], p *<* 10^*−*16^; IT: coefficient estimate = 0.003, 95% CI [0.001, 0.004], p *<* 0.001; Supp. Figure 4A). This indicates that iSTTC outperform ACF particularly in long spiking datasets with low IT values and high excitation strength, and on units with low firing rates (Figure 4B-C, right).

In line with these results, increasing the number of trials also strongly and log-linearly reduced the REE of iSTTC and PearsonR alike (Figure 4D; Supp. Figure 4B). Across the entirety of the parameter space, iSTTC outperformed PearsonR by an average of ~14% (coefficient estimate = −0.062, 95% CI [−0.068, −0.057], p *<* 10^*−*16^; Supp. Figure 4B) and consistently displayed a higher proportion of IT estimates within various ranges of the actual IT value (Figure 4E). The effect of IT estimation method interacted with the number of trials and the IT value (number of trials: coefficient estimate = 0.0098, 95% CI [0.005, 0.015], p *<* 10^*−*16^; IT: coefficient estimate = −0.015, 95% CI [−0.020, −0.01], p *<* 10^*−*16^; Supp. Figure 4B), indicating that it is particularly prominent at low trial number counts and high IT values (Figure 4D-E, right).

Lastly, iSTTC also converged faster than PearsonR to a stable IT estimate (Supp. Figure 4C-D). To quantify this, we resampled the same number of trials a varying number of times (50 to 1000), and assessed the consistency of the estimates. A larger fraction of iSTTC-derived IT values fell within various fixed ranges around the median, indicating greater stability (Supp. Fig. 4C). In addition, iSTTC showed a lower standard error of the median across all resampling iterations (Supp. Fig. 4D).

In summary, iSTTC consistently provides more accurate and stable IT estimates than traditional methods across varying data conditions, in particular for datasets with a low number of trials.

### 2.5 iSTTC improves inclusion rates and fit quality on epoched spiking activity

Current studies on ITs generally impose strict inclusion criteria that limit the analysis to a small percentage of recorded neurons. These criteria commonly include a minimal firing rate of 1 Hz, an absence of empty bins in the averaged binned spiking data, a minimal number of trials in epoched data, a declining ACF/ACF-like in a specific time window, an R-squared value *≥* 0.5, and manual inspection of fit quality. [1, 27, 7, 28, 29, 30]. Thus, we set out to investigate whether iSTTC allows the use of a larger proportion of single units for IT estimation, thereby increasing the representativeness and robustness of the measure. We quantified the proportion of rejected units due to a failed or negative R-squared fit, which indicates that the model performs worse than predicting the mean of the data. While IT estimations on continuous data led to a minimal proportion of rejected units (~0.25%), this proportion increased significantly when using the PearsonR method (~16.6%) and, to a much less extent, when using iSTTC on epoched data (~6.3%) (Figure 5A). The advantage conferred by iSTTC over PearsonR was not uniform throughout the simulation parameters. Rather, the usage of iSTTC reduced the failed fits particularly at low firing rates, high levels of excitation strength and low IT values (Figure 5B).

**Figure 5:**
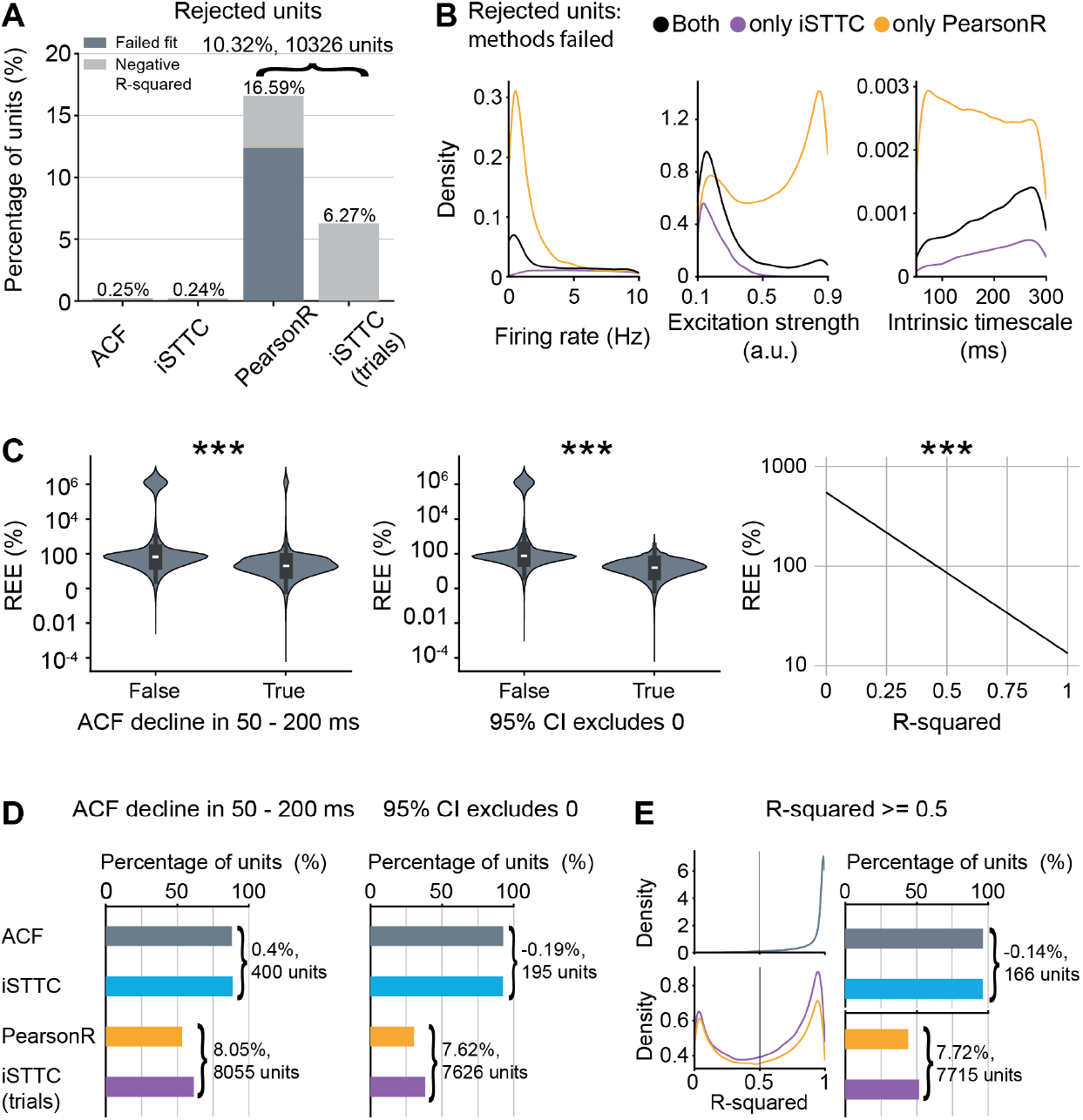
iSTTC allows inclusion of more units than PearsonR. **(A)** Bar plot displaying the percentage of excluded units across four methods. Color codes indicate exclusion reasons, with dark grey for failed exponential fits and light grey for negative R-squared values of the exponential fit (n = 23348 excluded fits across four methods, n = 248 excluded fits ACF, n = 238 excluded fits iSTTC, n = 16594 exluded fits PearsonR, n = 6268 excluded fits iSTTC (trials)). **(B)** Kernel density plot displaying the distribution of excluded units as a function of firing rate (left), excitation strength (middle), and IT (right) (n = 4469 units both methods failed, n = 12125 units only PersonR, n = 1799 units only iSTTC). **(C)** Violin plot displaying REE values for exponential fits where the autocorrelation function declined (vs. not declined) in the 50-200 ms range (left), and for fits where the 95% confidence interval of the estimated IT included vs. excluded zero (middle) (n = 376652 fits across four methods). Line plot displaying predicted REE values as a function of R-squared (right) (n = 376652 fits across four methods). Shaded areas represent 95% confidence intervals. Y-axes are plotted on a *log*_10_ scale. **(D)** Bar plots displaying the percentage of units with autocorrelation function decline in the 50–200 ms range (left, n = 291395 fits across four methods), and the percentage of units with 95% confidence intervals excluding zero (right, n = 253595 fits across four methods), across four methods. **(E)** Kernel density plot displaying the distribution of R-squared values (left), and bar plot displaying the percentage of units with R-squared *≥* 0.5 across four methods (right, n = 287967 fits across four methods). In **(C)**, left and middle, data are presented as median, 25th, 75th percentile, and interquartile range, with the shaded area representing the probability density distribution of the variable. In **(C)**, asterisks indicate a significant effect of the factor on REE. *** *p <* 0.001. Generalized linear models **(C)**.

To further assess whether commonly used quality criteria are valid indicators of estimation accuracy, we took advantage of the REE, which quantifies deviation from the IT ground truth value. We assessed the impact on REE of three quality parameters: the presence of an ACF/ACF-like decline in the 50 to 200 ms range, the R-squared value, and whether the 95% confidence interval of the IT estimate excludes zero. We found that each parameter was significantly predictive for REE (ACF/ACF-like decline: coefficient estimate = −0.946, 95% CI [−0.954, −0.937], p *<* 10^*−*16^; 95% CI excludes 0: coefficient estimate = −1.325, 95% CI [−1.332, −1.318], p *<* 10^*−*16^; R-squared: coefficient estimate = −1.613, 95% CI [−1.624, −1.602], p *<* 10^*−*16^, Figure 5C), suggesting that they are valid proxies of IT estimation quality. Importantly, since REE cannot be computed on experimental data where the ground truth is unknown, these results support their use to increase the reliability of IT estimates in practice.

Next, we compared how the proportion of units meeting each quality criterion varied across methods. Nearly all units showed an ACF/ACF-like decline in the 50 to 200 ms range in continuous data (~88%). This proportion dropped substantially in epoched data: 53.4% of units passed the criterion using PearsonR and 61.4% using iSTTC. Thus, iSTTC retained approximately ~8% more units than PearsonR (Figure 5D, left). A similar pattern was observed for the proportion of units whose IT estimate had a 95% CI that excluded zero. Here, ~92% of units met this criterion in continuous data, but only by 30.4% with PearsonR and 38% with iSTTC in epoched data, again highlighting an improvement of ~7.6% for iSTTC (Figure 5D, right). Finally, this trend persisted with the R-squared criterion: ~96% of units passed in continious data, compared to 43.9% with PearsonR and 51.6% with iSTTC in epoched data. Once more, iSTTC retained ~ 7.7% more units than PearsonR (Figure 5E).

To further examine how the amount of data influences the number of included units, we quantified the proportion of units meeting each quality criterion, as well as the proportion of excluded units, across increasing signal lengths and trial counts. In continuous data, both ACF and iSTTC showed high pass rates across all quality metrics, but iSTTC consistently resulted in fewer excluded units, particularly at shorter signal length (Supp. Fig. 5A). In epoched data, the difference between methods became more pronounced. Across all trial counts, iSTTC retained a higher percentage of units compared to PearsonR for each quality criterion and showed a lower percentage of excluded units (Supp. Fig. 5B). These results highlight that iSTTC is especially advantageous under challenging data conditions, such as short recordings or low trial numbers, where traditional methods are more likely to fail.

Taken together, these findings indicate that iSTTC allows a larger proportion of units to meet commonly used quality criteria, thereby increasing the yield and representativeness of IT estimates.

### 2.6 iSTTC outperforms traditional methods also on an experimental dataset

To assess the performance of iSTTC on continuous and epoched experimental data, we analyzed a subset of the Visual Coding Neuropixels dataset [35], [2]. The dataset comprised 5,775 single units from 26 mice (Figure 6A, Supp. Figure 6A). Recordings were obtained from six cortical areas: primary visual cortex (V1), lateromedial area (LM), anterolateral area (AL), rostrolateral area (RL), anteromedial area (AM), and posteromedial area (PM), and two thalamic areas: the lateral geniculate nucleus (LGN) and the lateral posterior nucleus (LP). To estimate baseline ITs, we used data from the Functional Connectivity sessions, which included 30 minutes of spontaneous activity during gray screen presentation. We then artificially created trials by randomly selecting 40 1s-long segments from each 30-minute spike train, following the same approach used in the synthetic dataset. Firing rates ranged from 0.06 to 69.7 Hz (median: 3.9 Hz), and local variance from 0.14 to 2.16 (median: 0.86) (Figure 6A, Supp. Figure 6A).

**Figure 6:**
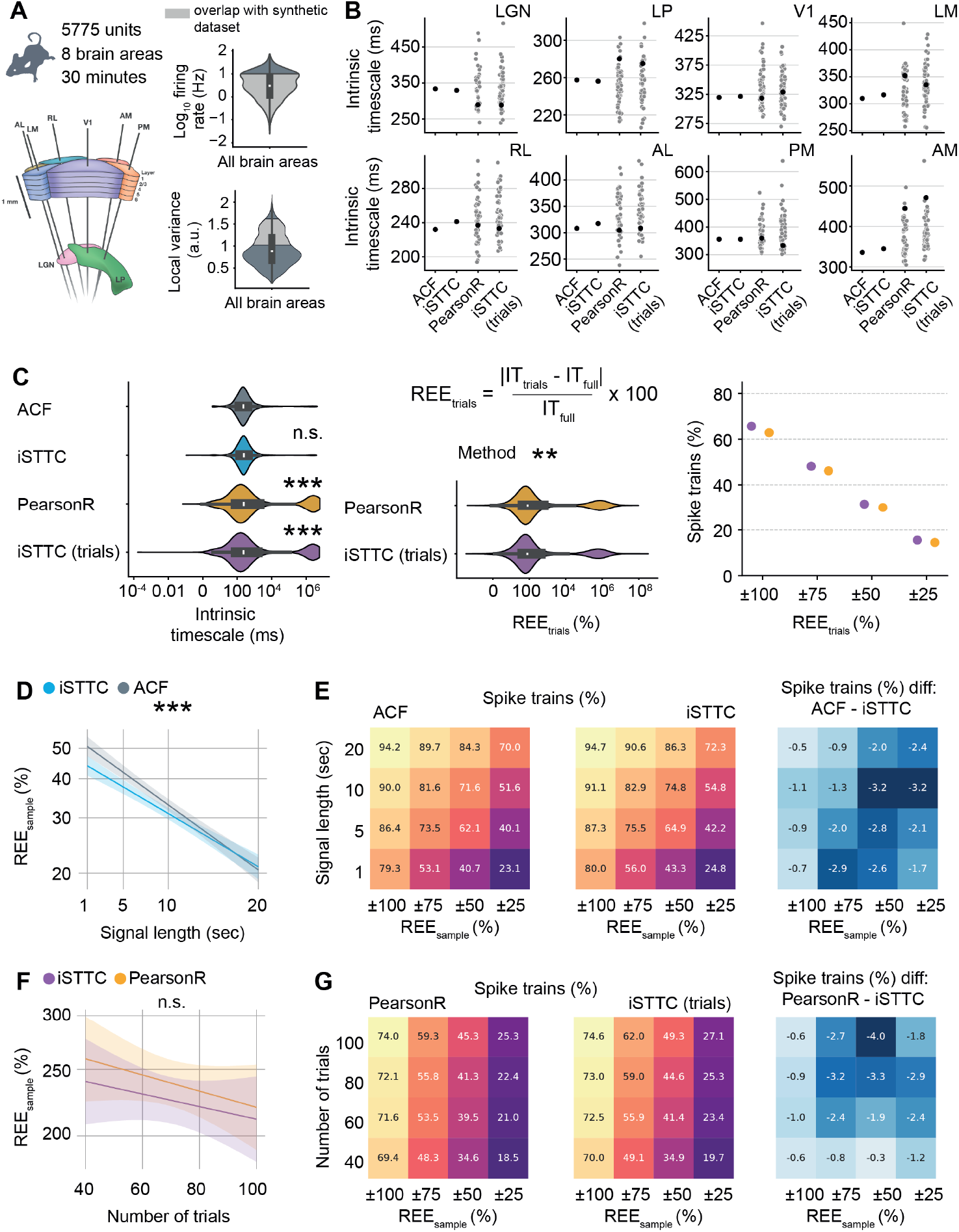
iSTTC outperforms ACF and PearsonR in IT estimation accuracy and robustness also on experimental data. **(A)** Schematic representation of Neuropixels recordings from the six visual cortical areas (V1, LM, RL, AL, PM, AM) and the two thalamic areas (LGN and LP), image source ([2]) (left), violin plots displaying the single units firing rate (top right) and local variation (bottom right). **(B)** Scatter plots displaying ITs at the brain area level. Black dots indicate area-level ITs used for the analyses in Supp. Figure 5B and 5C. Grey dots represent individual brain-area IT estimates for the trial-based methods across different sampling iterations (n = 50 samples). **(C)** Violin plots displaying the estimated ITs (n = 3053 single units, only units for which all methods produced an IT estimate are included) as a function of the estimation method (left). Violin plots displaying REE as a function of the estimation method (middle). Scatter plot displaying the percentage of spike trains with REE falling within progressively narrower bounds (right). **(D)** Line plot displaying predicted REE values for iSTTC and ACF as a function of signal length (n = 5674 single units per signal length, only units for which both methods produced an IT estimate for all signal lengths are included). Shaded areas represent 95% confidence intervals. Y-axes are plotted on a *log*_10_ scale. **(E)** Heatmap displaying the percentage of spike trains with REE within specific intervals for ACF (left) and iSTTC (middle) across varying signal lengths. Color codes for the proportion of spike trains, with warmer colors indicating higher percentages of spike trains. (Right) Heatmap displaying the difference between ACF and iSTTC. Negative values indicate better performance (lower REE) for iSTTC. Color codes for the magnitude of the difference. **(F)** Same as **(D)** for PearsonR and iSTTC (n = 4588 single units per number of trials, only units for which both methods produced an IT estimate for all numbers of trials are included). **(G)** Same as **(E)** for PearsonR and iSTTC. In **(A)** and **(C)** left and middle, data is presented as median, 25th, 75th percentile, and interquartile range, with the shaded area representing the probability density distribution of the variable.In **(C)**, asterisks indicate a significant effect of the method. In **(C)** left, ACF is used as a reference. In **(C)** middle, PearsonR is used as a reference. In **(D)**, asterisks indicate a significant effect of signal length. ** *p <* 0.01, *** *p <* 0.001. Generalized linear models with interactions **(C), (D)**, and **(F)**.

We estimated ITs at the brain area level using two continuous-signal methods and two trial-based methods. Full-signal methods yielded consistent estimates across areas with relatively narrow confidence intervals and a well-preserved hierarchical ordering of ITs, consistent with previous results [32] (Supp. Figure 6B-C). Conversely, trial-based methods produced more variable estimates, wider confidence intervals (Supp. Figure 6B) and inconsistent hierarchical ordering of ITs (Supp. Figure 6C).

To further assess the stability of trial-based estimates, we repeated the trial sampling procedure 50 times per unit and recomputed area-level ITs. Resampling led to high variability in the estimates across trial sets (Figure 6B; the black dots mark the values from the original sampling iteration used in Supp. Figure 6B-C). Together, these results highlight how the instability of trial-based methods also in experimental data can lead to inconsistent results and hierarchical ordering of ITs.

Next, we compared IT estimates at the single-unit level. ACF and iSTTC produced similar results with no significant differences between them (coefficient estimate = 0.030, 95% CI [−0.023, 0.085], p = 0.264), whereas trial-based methods differed significantly from ACF (PearsonR: coefficient estimate = 0.655, 95% CI [0.601, 0.709], p *<* 10^*−*16^; iSTTC trials: coefficient estimate = 0.624, 95% CI [0.570, 0.678], p *<* 10^*−*16^; Figure 6C left; Supp. Figure 6D). Importantly, when modeling the REEs we found that, on average, iSTTC yielded REEs that were ~15% smaller than PearsonR (coefficient estimate = −0.071, 95% CI [−0.117, −0.024], p = 0.0028; Figure 6C, middle; Supp. Figure 6E). Along the same lines, iSTTC led to a larger (~5%) proportion of IT estimates with an REE that was within a range of 100% to 25% of the IT estimated by iSTTC (Figure 6C, right).

We next quantified the effect of signal duration and trial number on IT estimated from continuous and epoched data, respectively. To this aim, we systematically varied the signal length (from 1 to 20 minutes) and the trial count (from 40 to 100 trials), and computed the REE with respect to the IT estimated using iSTTC on the full-signal. Similarly to synthetic data, increasing signal length led to a log-linear decrease in REE. Importantly, iSTTC outperformed ACF across the entirety of the dataset (~7% lower REEs, coefficient estimate = −0.031, 95% CI [−0.047, −0.016], p *<* 10^*−*5^; Supp. Figure 7A), an advantage that was particularly evident on short signals (Figure 6D; Supp. Figure 7A). Further, iSTTC consistently estimated a larger proportion of IT with an REE that fell within various ranges of the IT estimated on the full signal (Figure 6E).

On epoched data, increasing the number of trials also led to a log-linear decrease in REE, but no differences between iSTTC and PearsonR reached statistical significance (coefficient estimate = −0.025, 95% CI [−0.054, 0.004], p = 0.0894; Figure 6F, Supp. Figure 7B). However, REEs of ITs estimated with iSTTC consistently fell at a higher proportion within various ranges of the IT estimated on the full signal (Figure 6G). Moreover, iSTTC converged faster to an accurate IT estimation. To quantify this, we resampled the same dataset 50 to 1000 times with different and randomly selected trials, and assessed the consistency of the estimates. A larger fraction of iSTTC-derived IT values fell within various ranges around the median, indicating greater stability (Supp. Fig. 7C, left). In addition, iSTTC showed a lower standard error of the median across resampling iterations (Supp. Fig. 7D).

Lastly, we evaluated whether iSTTC allows to include a larger proportion of units also on experimental data. To this aim, we quantified the proportion of rejected units and the proportion of units meeting the same three quality criteria that we used for the synthetic data. In contrast to the synthetic dataset, we observed a high proportion of excluded units also in continuous data, with minimal differences between ACF and iSTTC (~21-22%). Similarly to our previous results, this proportion was substantially higher on epoched data. Also in this case, iSTTC outperformed PearsonR, yielding a rejection rate of 28.3% compared to 40.6%, a decrease of 12.4% (Supp. Fig. 8A). When comparing PearsonR and iSTTC on epoched data, iSTTC allowed to include a larger proportion of units with low firing rates and bursty firing patterns (Supp. Fig. 8B). In line with these results, on epoched data, iSTTC also retained a higher percentage of units passing each quality criterion, including ACF/ACF-like decline in 50 − 200 ms range (5.3% increase for iSTTC), 95% CI excludes 0 (3.6% increase for iSTTC), and R-squared *≥* 0.5 (4.7% increase for iSTTC) (Supp. Fig. 8C, D). On continuous data, iSTTC and ACF performed similarly (Supp. Figure 8C-D). These findings show that, also on epoched experimental data, iSTTC enables the use of a larger proportion of recorded units.

In conclusion, also on experimental data, iSTTC outperforms ACF and PearsonR IT estimation accuracy. Moreover, IT estimates based on continuous data are more robust and consistent than trial-based methods.

## 3 Discussion

In this study we introduce iSTTC, a novel method for accurately estimating ITs from SUA recordings. We demonstrate the advantages of iSTTC over currently used methods using synthetic and experimental data. Under a wide variety of conditions, iSTTC provides more accurate and stable IT estimates. Further, iSTTC accommodates both continuous and epoched data without bias, and generally allows for the inclusion of a larger number of single units. Together, these properties significantly enhance the accuracy, representativeness and robustness of IT estimations in neural circuits.

iSTTC is an extension of STTC [31], adapted specifically to address the challenges of IT estimation. Unlike traditional ACF approaches, iSTTC directly quantifies the autocorrelation-like function on binary spiking data without binning or smoothing. This reduces the number of arbitrary parameters required for the analysis. Crucially, iSTTC is also insensitive to 0-padding, and can therefore be seamlessly applied also to epoched data. When tested on synthetic data, iSTTC recovers ITs more accurately than other state-of-the-art methods, both on continuous and epoched signals. On continuous data, these advantages are especially pronounced under biologically realistic conditions, such as sparse firing and strong recurrent connectivity. Lastly, on epoched data, iSTTC produces a larger amount of ACF-like estimates that pass commonly used inclusion criteria, significantly expanding the proportion of neurons suitable for analysis. This increased single-unit inclusivity enhances statistical robustness and ensures IT estimates that better represent the underlying neural circuit. When investigating a state-of-the-art experimental dataset, iSTTC required shorter recording durations than traditional ACF methods to converge to stable solutions. On epoched experimental data, iSTTC remained more robust, converging more rapidly to estimates closer to those derived from continuous recordings. Further, we confirmed that substantially more neurons met inclusion criteria under iSTTC then PearsonR, thus producing more representative estimates of the IT of a given brain area.

An additional key insight from our analysis is the inherent instability in ITs estimated from epoched data, a common practice in the field. Our analyses on simulated and experimental data demonstrate that short trial lengths or limited trial numbers drastically reduce the reliability of IT measurements. Overall, estimating ITs on epoched data with a number and duration of trials that is similar to those employed in the literature [1, 27, 7, 34] leads to a roughly ten-fold increase in REEs when compared to IT estimated on continuous data. This warrants caution in interpreting ITs computed with this approach. While iSTTC mitigates this instability and improves estimation accuracy, we emphasize the importance of using either long uninterrupted recordings or, if epoched data is unavoidable, ensuring maximal trial length and number.

The methodological advances introduced by iSTTC have important implications for future research on ITs. Reliable estimation of ITs is critical for understating how they are regulated and to link them to cognitive and behavioral outcomes. Current theoretical frameworks suggest IT variability across brain areas underpins functional specialization [3]. By enabling more accurate and inclusive measurement of ITs, iSTTC provides a powerful tool to empirically test these theories across diverse neural systems. Moreover, given the wide heterogeneity of single-neuron ITs also within brain areas [7, 36, 15, 34], and the emerging evidence linking IT modulation to single neuron properties [3, 34], we anticipate that a robust metric which increases the proportion of neurons passing inclusion criteria like iSTTC will be particularly insightful to deepen our understanding of the principles governing ITs.

In conclusion, iSTTC expands the toolbox for IT estimation, overcomes several shortcomings of the current methods, and enhances both accuracy and representativeness of IT estimations. These improvements might lead to deeper insights into the mechanisms regulating ITs, ultimately bringing us closer to one of the fundamental goals of neuroscience: mapping behavioral actions onto neural circuits.

## 5 Author contributions

I.P., M.C., and I.L.H.-O. designed the study and wrote the paper. I.P. and M.C. developed the method and performed the analytical work. All authors approved the submitted version.

## 7 Materials and methods

### 7.1 Datasets

#### 7.1.1 Synthetic dataset

To generate synthetic spike trains with a specific intrinsic time constant, we implemented a univariate Hawkes point process with an exponential kernel, using Ogata’s thinning algorithm.

Hawkes process is defined to be a self-exciting temporal point process whose conditional intensity function *λ*(*t*) is defined as:

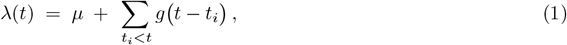

where *µ >* 0 is the baseline intensity and *g*(Δ), where Δ = *t − t*_*i*_, is an exponential excitation kernel defined as:

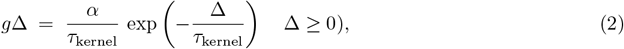

with *α ∈* [0, 1) governing the total integrated excitation (ensuring stability) and *τ* as the kernel’s time constant. Hence, explicitly:

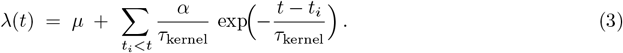

Because *α <* 1, the process remains subcritical and its long–term rate converges to 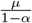.

For each spike train simulation, we specified four parameters: a target stationary firing rate *f* (in Hz), a desired time constant *τ* (in ms), an excitation strength *α* (dimensionless, with *α <* 1 for process stability), and a total simulation duration *d* (in ms).

The kernel time constant was computed as *τ*_kernel_ = *τ* (1 *− α*), and the baseline intensity *µ* in units of Hz was set to *µ* = *f* (1 *− α*).

The method is implemented in the *simulate hawkes thinning* function (available at https://github.com/iinnpp/isttc).

The generated datasets are described in the Table 3.

**Table 3:** Summary of synthetic datasets.

| Dataset | Parameters | Dataset file | Related figures |
| --- | --- | --- | --- |
| Parametric dataset | $f_{min} = 0.01$ Hz, $f_{max} = 10$ Hz,<br>$\alpha_{min} = 0.1$, $\alpha_{max} = 0.9$,<br>$\tau_{min} = 50$ ms, $\tau_{max} = 300$ ms,<br>$d = 600$ seconds,<br>number of spike trains = 100000.<br>Trial generation:<br>number of trials = 40,<br>trial duration = 1000 ms.<br>A total of 100 000 independent parameter sets were generated by sampling $f$, $\tau$, $\alpha$ uniformly within the predefined intervals. | spike_trains.pkl | Figure 2, Figure 3, Figure 4A, Figure 5 |
| Parametric datasets with a varying number of trials | For each spike train in the previously generated dataset (600 seconds in duration), four trial sets were created, each consisting of a different number of 1000 ms trials (40, 60, 80, and 100 trials). | trials40.pkl,<br>trials60.pkl,<br>trials80.pkl,<br>trials100.pkl | Figure 4B,C |
| Parametric datasets with a varying signal length | For each spike train in the previously generated dataset (600 seconds in duration), four truncated versions of lengths 60 seconds, 150 seconds, 300 seconds, and 450 seconds were generated. | length60.pkl,<br>length150.pkl,<br>length300.pkl,<br>length450.pkl | Figure 4D,E |

#### 7.1.2 Mouse dataset

We analyzed a subset of the Visual Coding Neuropixels dataset, which is publicly available via the Allen Brain Observatory [2, 35]. This dataset was previously used by [32] to calculate intrinsic timescales. This dataset comprises extracellular electrophysiological recordings from mouse brain acquired with Neuropixels probes. During the experiments, head-fixed mice were presented with a variety of visual stimuli. The dataset includes two sets from the two experimental pipelines, Functional Connectivity and Brain Observatory 1.1, which differ in their stimulus sequences. To compute baseline intrinsic timescales, we used data from the Functional Connectivity set, which contains a block of around 30 minutes of spontaneous activity recorded while the animals viewed a gray screen. For the analysis, we trimmed all recordings to 30 minutes. Recordings span six cortical areas (primary visual cortex (V1), lateromedial (LM), anterolateral (AL), rostrolateral (RL), anteromedial (AM), and posteromedial (PM)) and two thalamic areas (lateral geniculate nucleus (LGN) and lateral posterior nucleus (LP)). For analysis, we only included single units satisfying the following quality criteria (in agreement with [32]:

- presence ratio *≥* 0.9 (presence ratio 0.9 or higher means that the unit was present at least 90% of the recorded time),
- ISI (interspike interval) violations ≤ 0.5 (ISI violations *≤* 0.5 of lower means that contaminating spikes are occurring at most at roughly half the rate of “true” spikes for the unit),
- amplitude cutoff *≤* 0.01 (amplitude cutoff of 0.01 or lower indicates that at most 1% of spikes are missing from the unit).

After applying these criteria, 5775 units remained, with a minimum of n = 254 units (LGN) and n = 12 mice (LGN) and a maximum of n = 1063 units (V1) and n = 24 mice (V1 and AM) per brain area.

### 7.2 Autocorrelation function estimation

We used four methods to compute the autocorrelation function. Two were applied to the full spike train, the “classic” autocorrelation (ACF) on binned spike trains and iSTTC on non-binned spike trains. The other two were trial-based, using PearsonR on binned spike trains and iSTTC on concatenated non-binned spike trains.

#### 7.2.1 “Classic” autocorrelation - ACF

To compute the ACF on binned spiking data, we used the following equation:

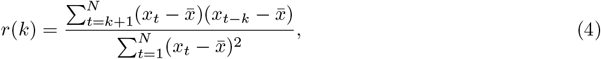

where *r*(*k*) is the autocorrelation at *k* lag and *N* is the total number of time points (bins) in the signal. We used the `statsmodels.tsa.stattools.acf` function for ACF calculation.

#### 7.2.2 i(ntrinsic)STTC

To compute the autocorrelation function on non binned spiking data, we used modified version of STTC (Spike time tilling coefficient) [31]. The STTC is defined as follows:

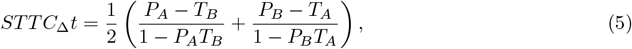

where *STTC*_Δ*t*_ is a correlation between spike trains *A* and *B* on the time lag Δ*t*; *T*_*A*_ (*T*_*B*_) is the proportion of total signal length that lies within *±* Δ*t* of any spike of *A* (*B*); *P*_*A*_ (*P*_*B*_) is the proportion of spikes from *A* (*B*) that lies within *±* Δ*t* of any spike of *B* (*A*).

The STTC formula is ill-suited for computing the autocorrelation. By design, STTC centers each time lag on individual spikes and uses expanding intervals around those spikes rather than a sliding window of variable size. As a result, applying STTC to the same spike train always yields a correlation of 1, since each spike coincides with itself.

To use the STTC for autocorrelation, we generated lagged versions of a spike train by shifting the original signal by a sequence of time lags. For each time lag *t*, for the spike train A, we truncated the original spike train by removing the spike times *< t* and for the spike train B truncated the end of the unshifted spike train by removing spike times *>*= *signal length − t* so that both sequences remained the same length. We then computed STTC between these two aligned truncated spike trains. Thus, a 50 ms shift yields the first autocorrelation lag, 100 ms the second, and so on. We defined the iSTTC autocorrelation at *k* lag as follows:

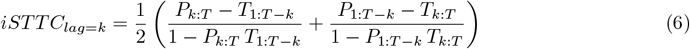

The iSTTC requires three parameters: the number of lags *n*_lags_, which specifies the total number of time lags over which the ACF is estimated; *lag shift*, which defines the increment between successive lags; and Δ*t*, within which the iSTTC evaluates spike train pair correlations. The method is implemented in the acf sttc function (available at https://github.com/iinnpp/isttc).

#### 7.2.3 PearsonR trial average

We implemented the trial average autocorrelation of spike counts (binned spike trains) using Pearson’s correlation coefficient as previously described [1, 27, 7, 28, 2].

The PearsonR trial average method requires as a parameter the number of lags *n*_lags_, which specifies the total number of time lags over which the ACF is estimated. An additional parameter is the *bin size*, which is used to bin the spike trains before applying PearsonR. The method is implemented in the *acf pearsonr trial avg* function (available at https://github.com/iinnpp/isttc).

Given a set of binned spiking data from multiple trials, represented as:

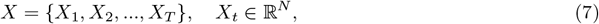

where *X*_*t*_ is the binned spikes time series data for trial *t*, and *N* is the number of time bins, the ACF for the first *n*_lags_ time lags was computed as follows:

**Step 1: Extract relevant time series** We define a subset of the time series that includes the first *n*_lags_ time bins across trials:

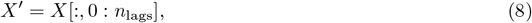

where *X*^*′*^ is a matrix of size *T × n*_lags_, with *T* trials and *n*_lags_ time bins.

**Step 2: Compute Pearson correlation for each lag** For each pair of time points *i* and *j*, where *j > i*, the Pearson correlation coefficient is computed as:

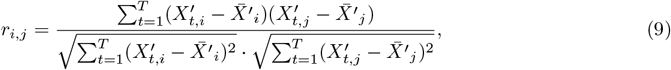

where *r*_*i,j*_ represents the Pearson correlation coefficient between time bins *i* and *j*, 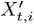 is the value of the time series at bin *i* for trial *t*, 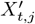 is the value of the time series at bin *j* for trial *t*, 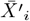 is the mean across trials for bin *i*, 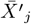 is the mean across trials for bin *j*.

These values are stored in an ACF matrix *R* of size *n*_lags_ *× n*_lags_, with:

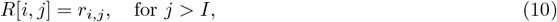

where the diagonal elements are set to *R*[*i, i*] = 1 (self-correlation at time lag 0).

**Step 3: Compute the trial average ACF** To obtain the trial average autocorrelation function, we compute the mean correlation for each lag *k*, considering the values along the diagonals of the ACF matrix:

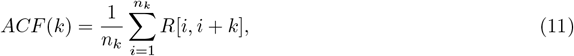

where *n*_*k*_ is the number of valid correlations available for each lag *k*.

The result is a one-dimensional autocorrelation function of length *n*_lags_.

#### 7.2.4 iSTTC (trials)

In the iSTTC for trials, all trials for a given unit are concatenated, with zero padding inserted between trials, to form a continuous spike-train signal. This concatenated signal is then used to compute the STTC in a manner analogous to iSTTC. However, the *T* term is calculated using the concatenated signal without zero padding.

The iSTTC for trials requires four parameters: the number of lags *n*_lags_, which specifies the total number of time lags over which the ACF is estimated; *lag shift*, which defines the increment between successive lags; Δ*t*, within which the STTC evaluates spike train pair correlations; and *zero_padding_len*, which is the length of zero padding appended to each concatenated trial. The method is implemented in the `acf_sttc_trial_concat` function (available at https://github.com/iinnpp/isttc).

### 7.3 Intrinsic timescale estimation

To estimate the intrinsic time constant *τ* from the signal autocorrelation function, we fit the function to a single-exponential function defined as:

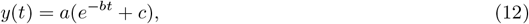

where *a* is the amplitude, *b* is the decay rate, and *c* is an offset parameter. The time constant *τ*, which characterizes the decay rate, is given by:

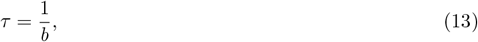

We estimate the parameters *a, b, c* using non-linear least squares fitting with the `curve_fit` function from SciPy. To ensure numerical stability, we constrain *b* to be strictly positive during fitting: *b >* 0.

We estimated confidence interval for *tau* as following: since *τ* depends on *b*, its standard error is derived using error propagation:

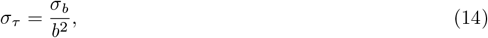

where *σ*_*b*_ is the standard error of *b*, obtained from the covariance matrix of the fitted parameters.

To compute the 95% confidence interval for *τ*, we use the Student’s t-distribution, which accounts for small sample sizes:

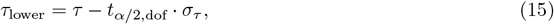

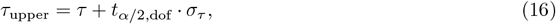

where:

- *t*_*α/*2,dof_ is the critical t-value for a 95% confidence level,
- dof is the degrees of freedom (number of data points *−* number of parameters).

### 7.4 Intrinsic timescale estimation in the synthetic dataset

Intrinsic timescales were estimated at the level of individual units using four methods. The autocorrelation function was computed for all units without any exclusion criteria. If the computation failed at a specific time lag, the corresponding value was set to `NaN`.

For iSTTC, the following parameters were used: number of lags *n*_lags_ = 20, *lag_shift* = 50 ms and Δ*t* = 25 ms. When applying iSTTC to trial-based data, we additionally used a *zero_padding_len* = 3000 ms, which corresponds to the amount of zero-padding appended to each concatenated trial. For ACF and PearsonR, spike trains were binned with a bin size of 50 ms, and the number of lags was set to *n*_lags_ = 20. The intrinsic timescale was determined by fitting a single-exponential decay function to the autocorrelation curve, starting from a time lag of 1. Autocorrelation functions containing NaN values were excluded from the fitting process.

### 7.5 Intrinsic timescale estimation in the mouse dataset

Intrinsic timescales were estimated at both the single-unit and brain-area levels using the same four methods. The autocorrelation function was computed for all units without any exclusion criteria. If the computation failed at a specific time lag, the corresponding value was set to `NaN`.

The same parameter settings were used as in the synthetic dataset.

For single units, the intrinsic timescale was determined by fitting a single-exponential decay function to the autocorrelation curve, starting from a time lag of 1. For brain areas, the fit was performed starting from a time lag of 2. Autocorrelation functions containing NaN values were excluded from the fitting process. To estimate the area-level intrinsic timescale, all valid ACFs (those without NaNs) from the area were used to fit a single exponential function.

### 7.6 Local variation

Local variation *Lv* was computed as follows [33]:

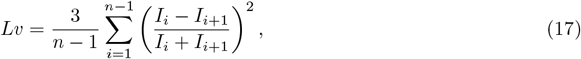

where *I*_*i*_ and *I*_*i*+1_ are the *i*-th and *i*+1st ISIs, and n is the number of ISIs.

Lv equals 0 for regular spike trains, 1 for Poisson spike trains and is above 1 for bursty spike trains.

### 7.7 Relative estimation error

Relative estimation error *REE* was computed for the synthetic dataset as follows:

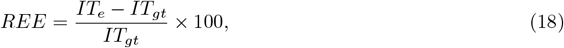

where *IT*_*e*_ is an estimated intrinsic timescale and *IT*_*gt*_ is the ground truth intrinsic timescale.

To compute the REE of PearsonR and iSTTC trials in the mice dataset, we used as ground truth the intrinsic timescale estimated using ACF or iSTTC, respectively. For the REE of methods applied to spike trains of varying lengths or different numbers of trials, the intrinsic timescale computed from the full signal was used as ground truth.

### 7.8 Statistical analysis

Statistical analysis was done in R. Non-nested data were analyzed with (generalized) linear models and nested data with (generalized) linear mixed-effects models. Initial model selection was based on exploratory analysis of the data, and then the goodness of fit was evaluated using explained variance (R-squared) and residuals distribution (check_model function from performance R package). When several models were fit on the same data, the model fit was compared using compare performance function from performance R package. 95% confidence intervals were computed using confint R function, p-values for linear mixed-effects models were computed with the lmerTest R package, post hoc analysis with Tukey multiple comparison corrections was done using emmeans R package, and the model fit was plotted using sjPlot and ggplot2 R packages.

### 7.9 Code availability

The complete analysis pipeline, including synthetic dataset and figures generation, is available at the following open-access repository: https://github.com/iinnpp/isttc.

### 7.10 Data availability

The synthetic dataset generated in this study is available at https://gin.g-node.org/iinnpp/isttc. The experimental dataset is available via the Allen Brain Observatory at https://allensdk.readthedocs.io/en/latest/visual_coding_neuropixels.html.

## 8 Supplementary information

**Supplementary Figure 1.**
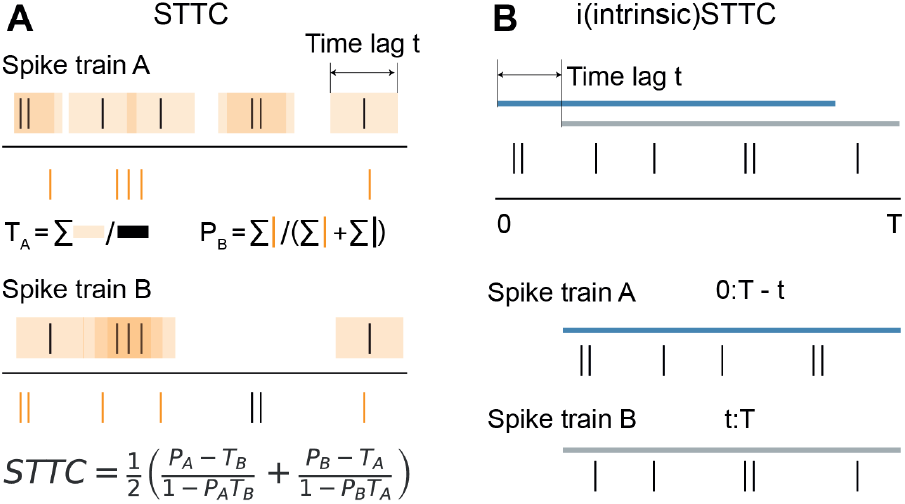
Schematic representation of spike time tiling coefficient calculation (STTC) and its adaptation for intrinsic timescale estimation (iSTTC). **(A)** Schematic representation of STTC calculation, modified from [31] and [37]. STTC quantifies the correlation between spike trains A and B at a time lag of t. *T*_*A*_ (*T*_*B*_) denotes the proportion of the signal within t/2 of any spike in A (B), and *P*_*A*_ (*P*_*B*_) the proportion of spikes in A (B) that fall within t/2 of a spike in B (A). **(B)** Schematic illustration of iSTTC. Spike trains A and B are created by temporally shifting and truncating the original spike train, after which the standard STTC formula is applied as in **(A)**.

**Supplementary Figure 2.**
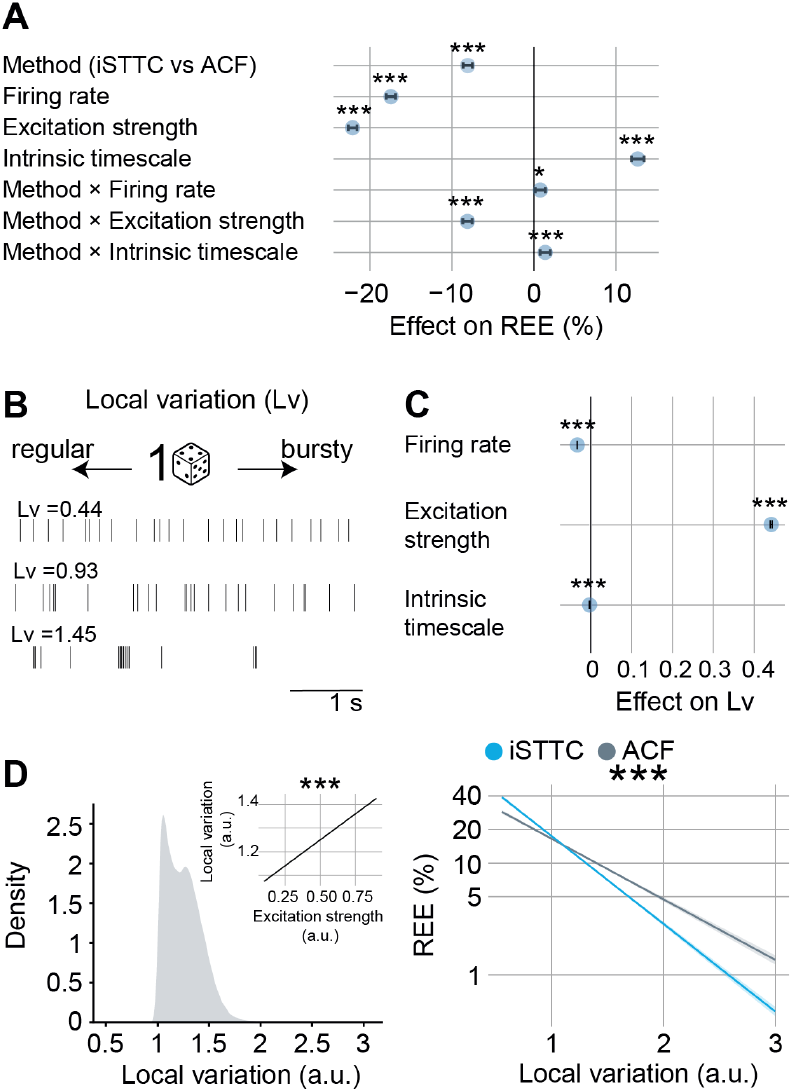
Local variation is an experimental proxy for the excitation strength parameter of the Hawkes point process. **(A)** Effect plot of the statistical model used to investigate the REEs plotted in Figure 2C-D. Negative values indicate reduced REE. ACF is used as the reference method. The x symbol denotes an interaction term between factors. **(B)** Schematic representation of local variation (Lv), modified from [33], and examples of spike trains with regular, random, and bursty firing patterns. **(C)** Effect plot of the statistical model used to investigate how Lv correlates with the Hawkes point parameters. **(D)** KDE plot displaying the distribution of Lv in the synthetic dataset, and, in the inset, the line plot displaying predicted Lv values across excitation strength (left). Line plot displaying predicted REE values for iSTTC and ACF across Lv (n = 10^5^ single units) (right). Shaded areas represent 95% confidence intervals. Y-axes are plotted on a *log*_10_ scale. In **(A)** and **(C)**, data is presented as mean difference (blue dots) with 95% confidence interval. In **(A), (C)**, and **(D)**, asterisks indicate a significant effect. * *p <* 0.05, *** *p <* 0.001. Generalized linear models with interactions **(A)** and **(D)**. Generalized linear model **(C)**.

**Supplementary Figure 3.**
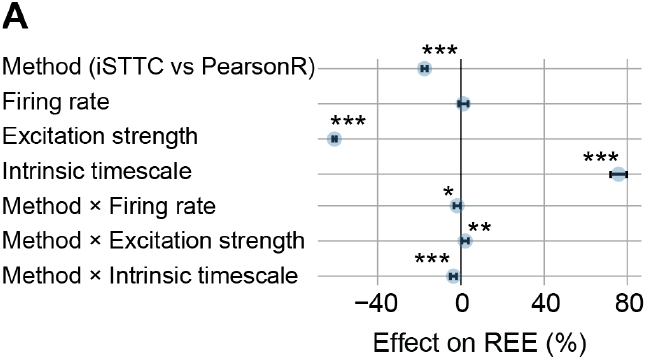
iSTTC yields significantly lower REEs than PearsonR on epoched data. **(A)** Effect plot of the statistical model used to investigate the REEs plotted in Figure 3B-C. Negative values indicate reduced REE, and PearsonR is used as the reference method. The x symbol denotes interaction terms between factors. In **(A)**, data is presented as mean differences (blue dots) with 95% confidence intervals. In **(A)**, asterisks indicate a significant effect. * *p <* 0.05, ** *p <* 0.01, *** *p <* 0.001. Generalized linear model with interactions **(A)**.

**Supplementary Figure 4.**
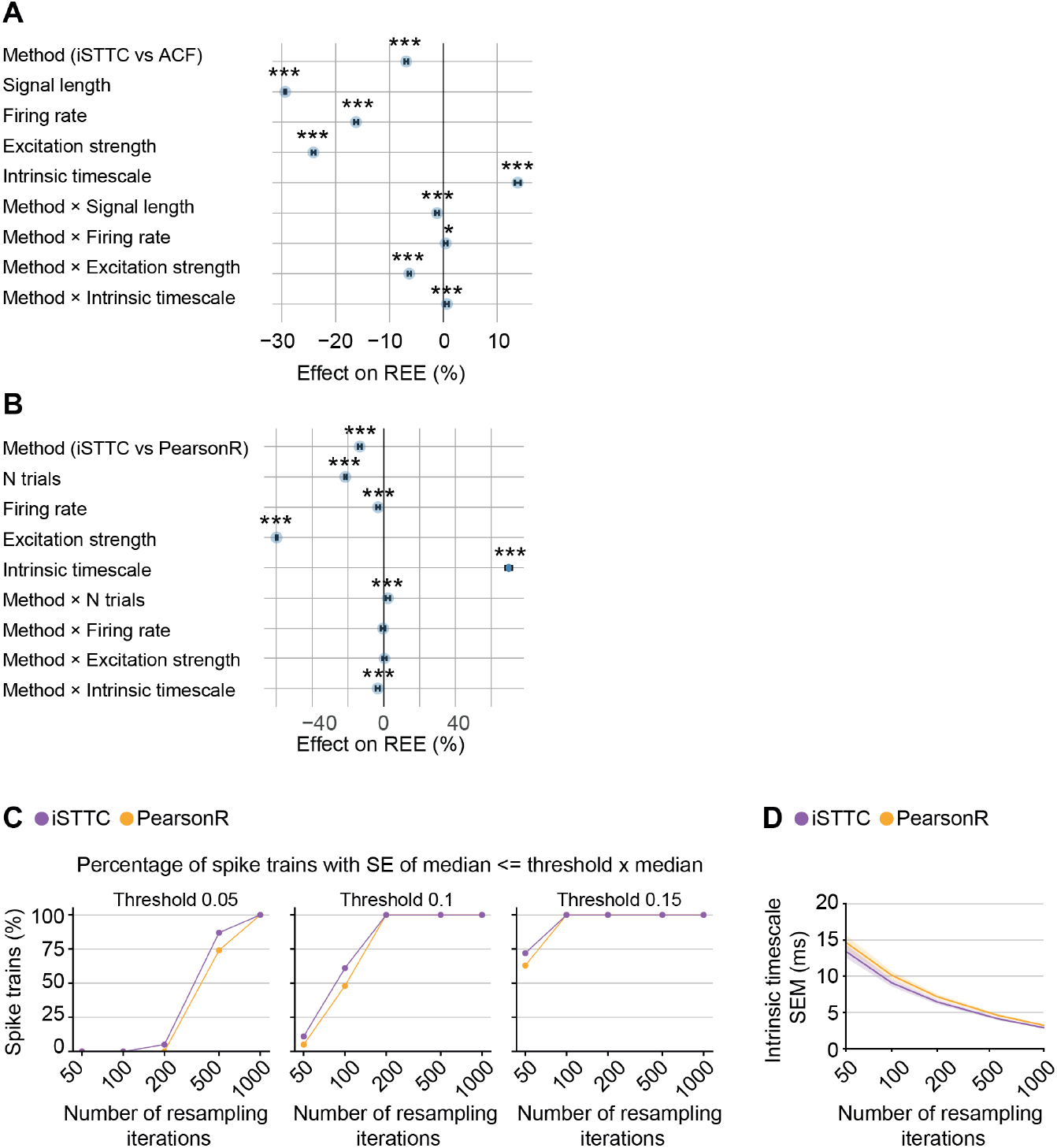
Epoched and short continuous data lead to unreliable IT estimates. **(A)** Effect plot of the statistical model used to investigate the REEs plotted in Figure 4A. Negative values indicate reduced REE, and ACF is used as the reference method. The x symbol denotes interaction terms between factors. **(B)** Same as **(A)** for the REEs plotted in Figure 4B. **(C)** Line plot showing the percentage of spike trains whose standard error of the median IT estimate falls within a specified tolerance range (5% to 15%) across resampling iterations. **(D)** Line plot displaying the IT estimate standard error of the median as a function of the number of resampling iterations. In **(A)** and **(B)**, data is presented as mean differences (blue dots) with 95% confidence intervals. In **(D)**, data is presented as mean with 95% confidence intervals. In **(A)** and **(B)**, asterisks indicate a significant effect. * *p <* 0.05, *** *p <* 0.001. Generalized linear model with interactions **(A)** and **(B)**.

**Supplementary Figure 5.**
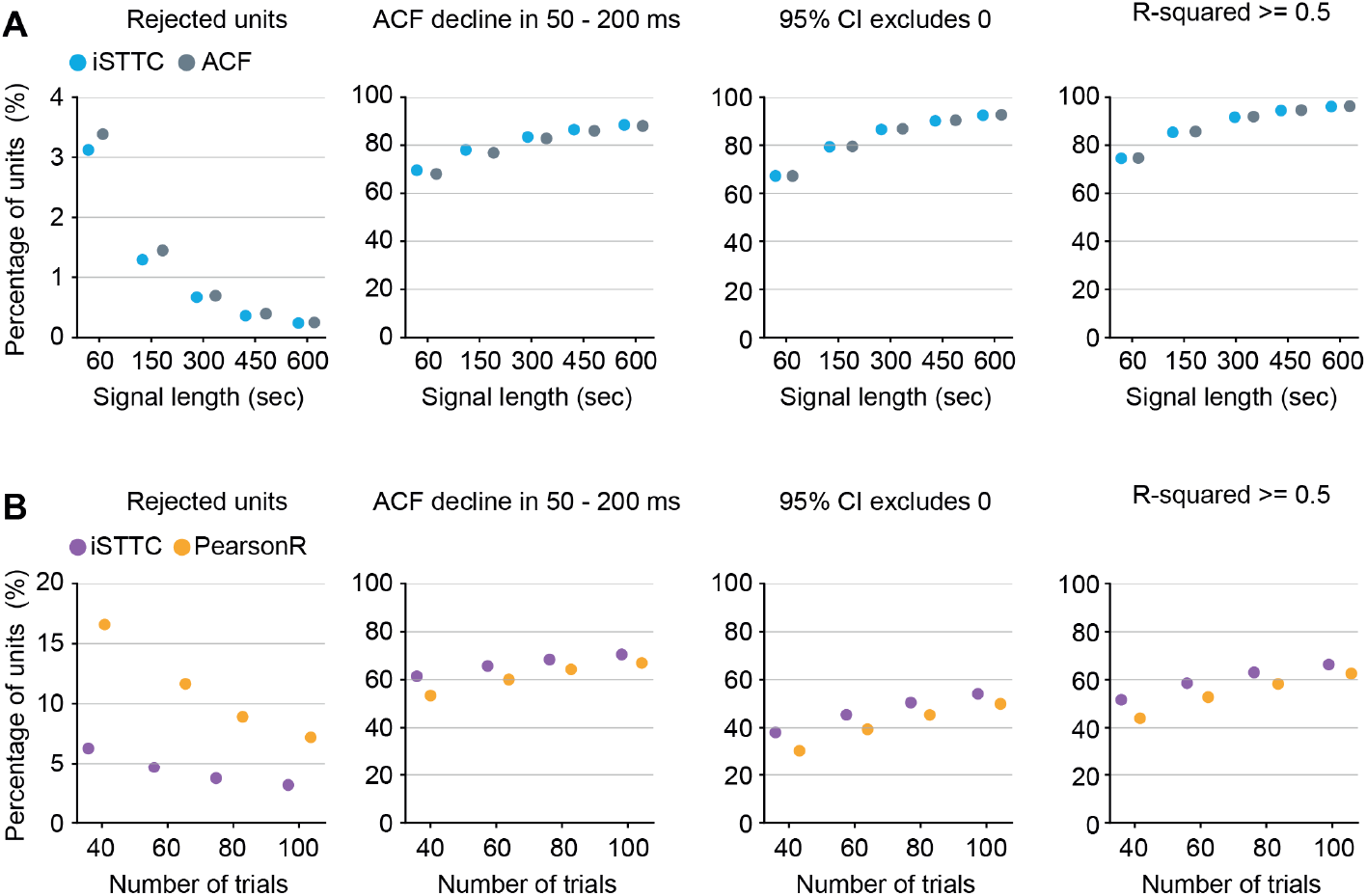
iSTTC allows inclusion of more units than PearsonR or ACF, especially for low trial counts and short signals. **(A)** Scatter plots showing the percentage of units across increasing signal lengths for ACF and iSTTC. From left to right: percentage of excluded fits, percentage of units with autocorrelation function decline in the 50–200 ms range, percentage of units with 95% confidence intervals excluding zero, and percentage of units with R-squared *≥*0.5. **(B)** Same as **(A)**, but for PearsonR and iSTTC across increasing numbers of trials.

**Supplementary Figure 6.**
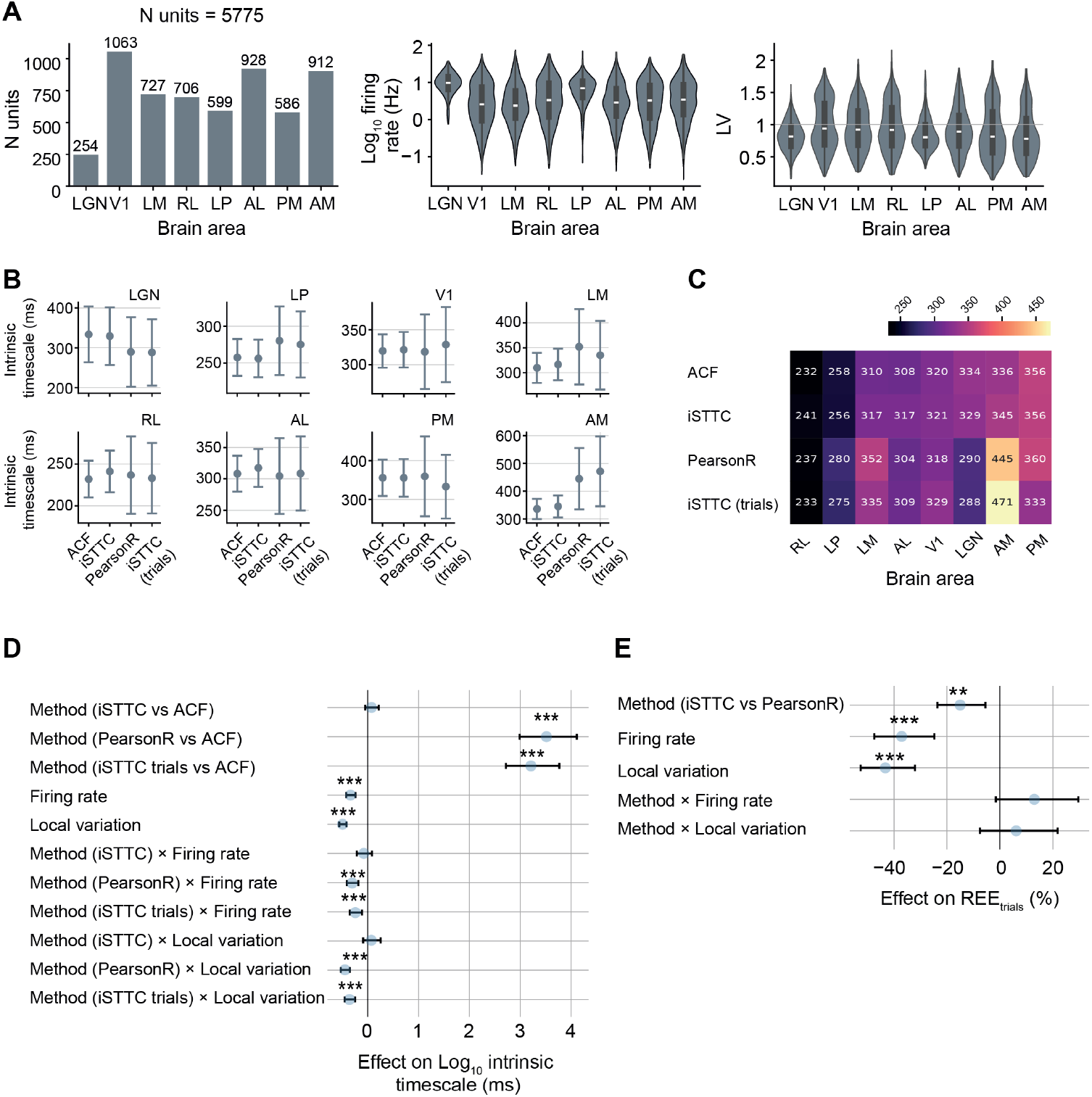
Trial-based IT estimation methods yield unreliable IT estimates on an experimental dataset. **(A)** Bar plot displaying the number of units per brain area (left), violin plot displaying the firing rate per brain area (middle), and violin plot displaying the Lv per brain area (right). The grey line denotes the Lv = 1 that corresponds to a random firing pattern. **(B)** Scatter plot displaying brain area level ITs estimated by four methods across individual brain areas. **(C)** Heatmap displaying brain area level ITs estimated by four methods across individual brain areas. Color codes for the estimated IT, with warmer colors indicating higher ITs. **(D)** Effect plot of the statistical model used to investigate the ITs plotted in Figure 5C, left. Negative values indicate reduced REE, and ACF is used as the reference method. The x symbol denotes interaction terms between factors. **(E)** Same as **(D)** for the REEs plotted in figure 5C, middle. In **(A)**, data is presented as median, 25th, 75th percentile, and interquartile range, with the shaded area representing the probability density distribution of the variable. In **(B)**, data is presented as mean with 95% confidence intervals. In **(D)** and **(E)**, data is presented as mean differences (blue dots) with 95% confidence intervals. In **(D)** and **(E)**, asterisks indicate a significant effect. ** *p <* 0.01, *** *p <* 0.001. Generalized linear model with interactions **(D)** and **(E)**.

**Supplementary Figure 7.**
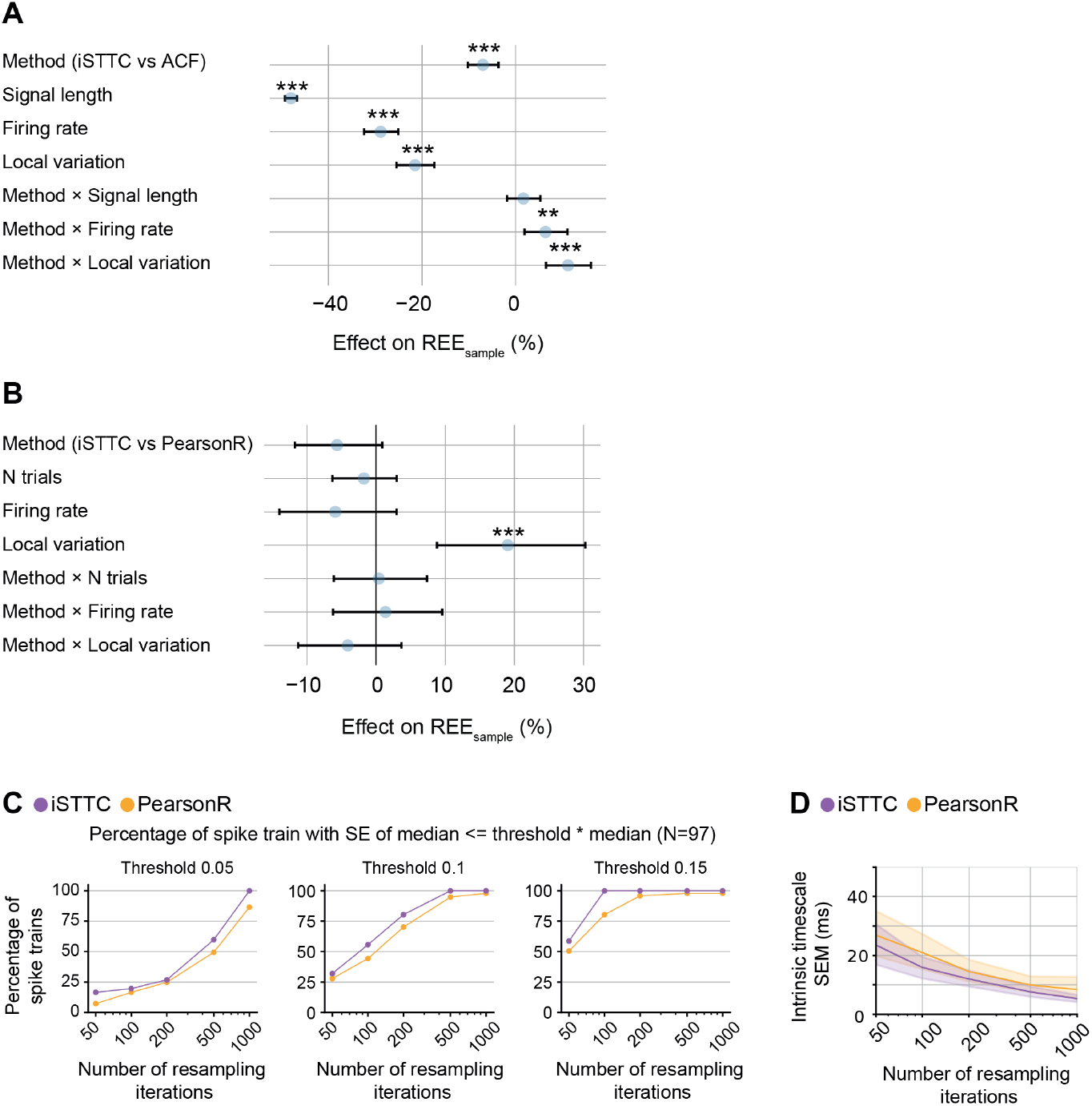
iSTTC outperforms ACF and PearsonR on short continuous and epoched data. **(A)** Effect plot of the statistical model used to investigate the REEs plotted in Figure 5D. Negative values indicate reduced REE, and ACF is used as the reference method. The x symbol denotes interaction terms between factors. **(B)** Same as **(A)** for the REEs plotted in Figure 5F. **(C)** Line plot showing the percentage of spike trains whose standard error of the median IT estimate falls within a specified tolerance range (5% to 15%) across resampling iterations. **(D)**Line plot displaying the IT estimate standard error of the median as a function of the number of resampling iterations. In **(A)** and **(B)**, data is presented as mean differences (blue dots) with 95% confidence intervals. In **(D)**, data is presented as mean with 95% confidence intervals. In **(A)** and **(B)**, asterisks indicate a significant effect. *** *p <* 0.001. Generalized linear model with interactions **(A)** and **(B)**.

**Supplementary Figure 8.**
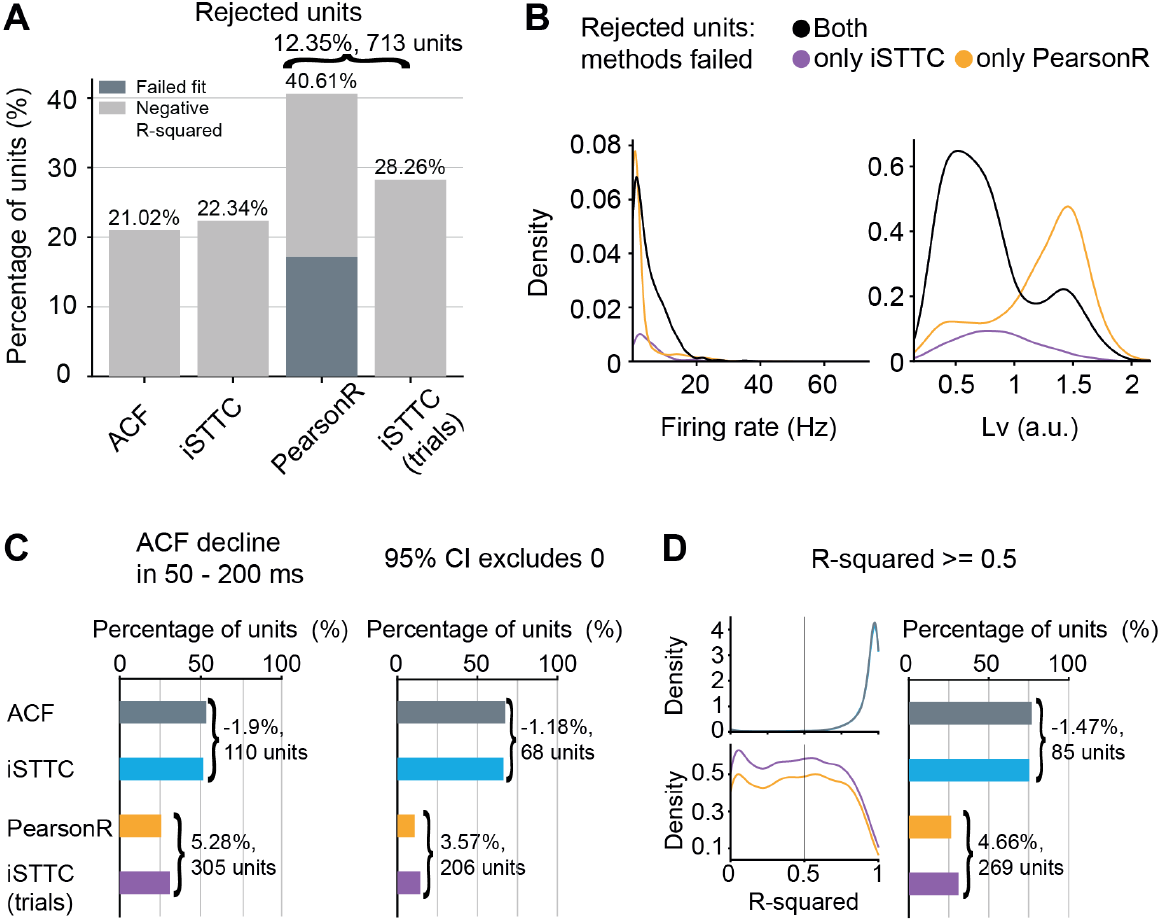
iSTTC allows inclusion of more units than PearsonR also on experimental dataset. **(A)** Bar plot displaying the percentage of excluded units across four methods. Color codes indicate exclusion reasons, with dark grey for failed exponential fits and light grey for negative R-squared values of the exponential fit (n = 6481 excluded fits across four methods, n = 1214 excluded fits ACF, n = 1290 excluded fits iSTTC, n = 2345 exluded fits PearsonR, n = 1632 excluded fits iSTTC (trials)). **(B)** Kernel density plot displaying the distribution of excluded units as a function of firing rate (left) and local variation Lv (right) (n = 1412 units both methods failed, n = 933 units only PersonR, n = 220 units only iSTTC). **(C)** Bar plots displaying the percentage of units with autocorrelation function decline in the 50–200 ms range (left, n = 9331 fits across four methods), and the percentage of units with 95% confidence intervals excluding zero (right, n = 9146 fits across four methods), across four methods. **(E)** Kernel density plot displaying the distribution of R-squared values (left), and bar plot displaying the percentage of units with R-squared *≥* 0.5 across four methods (right, n = 12102 fits across four methods).

